# A chromosome-level genome of the king penguin (*Aptenodytes patagonicus*): a resource for linking genotype to fitness in a long-lived vertebrate

**DOI:** 10.1101/2025.03.13.642884

**Authors:** Josephine R. Paris, Flávia A. Nitta Fernandes, Camilla A. Santos, Damon-Lee B. Pointon, Jonathan M. D. Wood, Joan Ferrer Obiol, Judit Salces-Ortiz, Rosa Fernández, Robin Cristofari, Céline Le Bohec, Emiliano Trucchi

## Abstract

The king penguin (*Aptenodytes patagonicus*) is a long-lived seabird of the Southern Ocean that is emerging as a promising model in the wild for studying evolution. We present a high-quality, haplotype-resolved 1.35 Gb chromosome-level genome of an adult female king penguin - ‘Pen/Se-guin’ - assembled using PacBio HiFi long-reads and Hi-C proximity data. A total of 95% of the assembly is assigned to 34 chromosomes (32 autosomes, plus the Z and W chromosomes). The primary haplotype has a BUSCO completeness of 97.2%, and across both haplotypes the assembly has a *k*-mer completeness of 99.7% and a quality value of 63.8. We also assembled a circularised mitogenome (20,520 bp) containing the avian tandem duplication. Repetitive sequence annotation showed that 16.3% of the genome comprises repetitive elements, with LINEs representing the most abundant transposable element class (5.6%). Gene prediction using an extensive multi-tissue RNA-seq dataset resulted in 18,081 predicted protein-coding genes, of which 17,081 were functionally annotated, with a BUSCO completeness of 98.4% and an OMArk completeness of 97.3%. Compared with the previous draft genome, the presented genome shows a 30-fold increase in assembly contiguity and a substantially improved genome annotation, exceeding the standards of the Earth BioGenome Project and Vertebrate Genomes Project. This genome provides a foundation for ongoing and future studies using the king penguin to investigate genotype–fitness relationships, ageing, life-history evolution, and adaptation.

## Introduction

The king penguin (*Aptenodytes patagonicus*) is a seabird of the Southern Ocean that has been the focus of extensive ecological and evolutionary research over the past several decades (e.g. Stonehouse, 1960; Oberlin et al. 2025). Long-term monitoring of individuals has generated extensive behavioural, demographic, and physiological data, providing unique opportunities for investigating evolutionary questions in the wild (e.g. Van Heezik et al., 1994; Le Vaillant et al., 2016; Bardon et al., 2026; Cristofari et al., 2026). However, linking genotype to phenotype and fitness has remained challenging due to the absence of a high-quality reference genome. The current king penguin assembly (GCA_010087175.1; Pan et al., 2019) is highly fragmented, comprising 15,968 scaffolds with an N50 of only 2.9 Mb and an incomplete annotation. More broadly, chromosome-level genomes remain scarce across penguins, with only two currently available: the Humboldt penguin (*Spheniscus humboldti*; GCA_027474245.1) and African penguin (*Spheniscus demersus*; GCA_050947425.1). To establish the king penguin as a model for evolutionary studies and to advance comparative genomics in penguins more broadly, we generated a high-quality chromosome-level genome assembly, including a mitogenome and comprehensive annotation for the species.

King penguins breed on subantarctic islands across the Southern Ocean, from 46°S at the Prince Edward and Crozet Islands to 54°S on South Georgia and Macquarie Island and have recently expanded their breeding range to the Falkland Islands (51°S) and Tierra del Fuego (53°S) (Pistorius et al., 2012; Gherardi-Fuentes et al., 2019). The species exhibits one of the longest breeding cycles of any bird, extending over 14 months (Descamps et al., 2002), and is long-lived, with an average lifespan of approximately 20 years in the wild (Cristofari et al., 2026). Combined with decades of individual-based monitoring, these characteristics make it particularly well suited for studying the ecological and evolutionary determinants of survival and reproduction.

King penguins are also an important model for understanding biological responses to environmental change (Le Bohec et al., 2008; Bardon et al., 2026). Their foraging ecology is tightly coupled to the Antarctic Polar Front, whose projected shifts are expected to drive major population declines or colony relocations during the coming century (Bost et al., 2015; Cristofari et al., 2018), while their large effective population size and weak population structure make the species informative for studying adaptive responses to rapid environmental change (Trucchi et al., 2014, 2025).

Here, we present ‘Pen/Se-guin’, a high-quality chromosome-level genome assembly for the king penguin. We name the genome after Louise Séguin, one of the first recorded European women known to travel to the Antarctic region. This assembly provides a valuable resource for investigating the genomic, transcriptomic, and epigenomic basis of fitness in the wild, while also supporting comparative genomic studies of penguin responses to past and future environmental change.

## Methods

### Biological materials

Samples for genome assembly were collected in 2022 from a 21-year-old female king penguin from *La Baie Du Marin* colony on Possession Island, Crozet Archipelago (46.43°S, 51.86°E). The individual was captured while leaving the colony during the breeding season. Blood aliquots (approx. 100 µl) were collected from the brachial vein using a heparinised syringe. Samples were preserved in 95% ethanol at −80°C and were shipped to the CNRS/Université de Strasbourg, France. One aliquot was used for PacBio long-read sequencing library preparation, and another for Hi-C library preparation. Tissues used for RNA-seq (brain, liver, kidney, pectoral muscle, and skin) were collected from freshly predated chick carcasses from the same colony (see Paris et al., 2025 for details). All procedures employed during fieldwork were approved by the French Ethical Committee and the French Polar Environmental Committee, and authorisations to enter the breeding site and to collect samples were delivered by the “Terres Australes et Antarctiques Françaises” (permit nos 2015-52, 2015-105, 2021-51, 2021-151).

### DNA extraction, library preparation, and sequencing

DNA extraction, library preparation and sequencing of the PacBio data were performed at BGI Group (Shenzhen, China). Genomic DNA was extracted from whole blood using a column-based protocol: the sample was lysed in preheated animal lysis buffer with proteinase K and RNase A (55 °C, 2 h), and the cleared lysate purified on a gravity-flow anion-exchange column, followed by isopropanol precipitation, a 75% ethanol wash and resuspension in elution buffer. SMRTbell libraries were prepared following the manufacturer’s standard workflow: genomic DNA was sheared, single-stranded overhangs were removed, and DNA damage repair, end repair and A-tailing were performed, before ligation of hairpin adapters and size selection. The library was sequenced on a PacBio Revio instrument (Pacific Biosciences, Menlo Park, CA, USA). An initial run produced 38.42 Gb of HiFi data and an additional SMRT Cell a further 20.12 Gb, giving 3,976,812 HiFi reads totalling 58.54 Gb (read length N50 = 15,051 bp; mean read length = 14,720 bp; 95.7% of bases ≥Q20).

Hi-C libraries were prepared using the Hi-C High-coverage kit (Arima Genomics) at the Metazoa Phylogenomics and Genome Evolution Lab (Institute of Evolutionary Biology - CSIC-UPF). Sample concentration was assessed using a Qubit DNA HS Assay kit (Thermo Fisher Scientific) and library preparation was carried out with the ACCEL-NGS® 2S PLUS DNA LIBRARY KIT (Swift Bioscience) using 2S Set A single indexes (Swift Biosciences). Library amplification was carried out with the KAPA HiFi DNA polymerase (Roche). Hi-C libraries were sequenced on an Illumina NovaSeq 6000 to produce 126 Gbp of paired-end read data. Hi-C data were cleaned with fastp v0.20 (Chen et al., 2018) using a quality threshold of 20, discarding reads <75 bp.

### Genome assembly

Software versions and parameters are given in Table 1. *K*-mer counting was performed on 21-mers derived from the HiFi data using DSK (Rizk et al., 2013). Resulting *k*-mer counts were used as input to GenomeScope2 (Ranallo-Benavidez et al., 2020) to estimate the haploid genome size, repetitiveness, and heterozygosity. The Pacbio HiFi data and Hi-C data were used to perform a Hi-C integrated assembly using Hifiasm (Cheng et al., 2021). Scaffolding was performed using YaHS (Zhou et al., 2023), with Hi-C reads aligned using Chromap (Zhang et al., 2021). Scaffolded assemblies for both haplotypes were used as input to the Nextflow nf-core pipeline (Ewels et al., 2020) curationpretext (pipelines.tol.sanger.ac.uk/curationpretext; Pointon, 2025) to generate Hi-C contact maps for manual curation (Howe et al., 2021). Manual curation of the combined haplotypes was performed using PretextView (github.com/sanger-tol/PretextView) with the purpose of correcting misassemblies, identifying sex chromosomes, and achieving a resolved chromosome-level assembly for the species. Identification of scaffolds belonging to the sex chromosomes was performed using merged map BUSCO plots using *Gallus gallus* BUSCO genes. Scaffolds of potential microchromosome scaffolds were identified using MicroFinder (github.com/sanger-tol/MicroFinder; Mathers et al., 2026), which were visualised and assembled based on synteny dot plots (at dot.sandbox.bio) generated via nucmer alignments in MUMmer4 (Marçais et al., 2018) against five avian species (Supplementary Table 1). The AGP PretextView output file was used as input for the pretext-to-asm.py script and the genome curated fasta files for each haplotype were generated using the multi_join.py script, available at github.com/sanger-tol/rapid-curation.

**Table 1.**
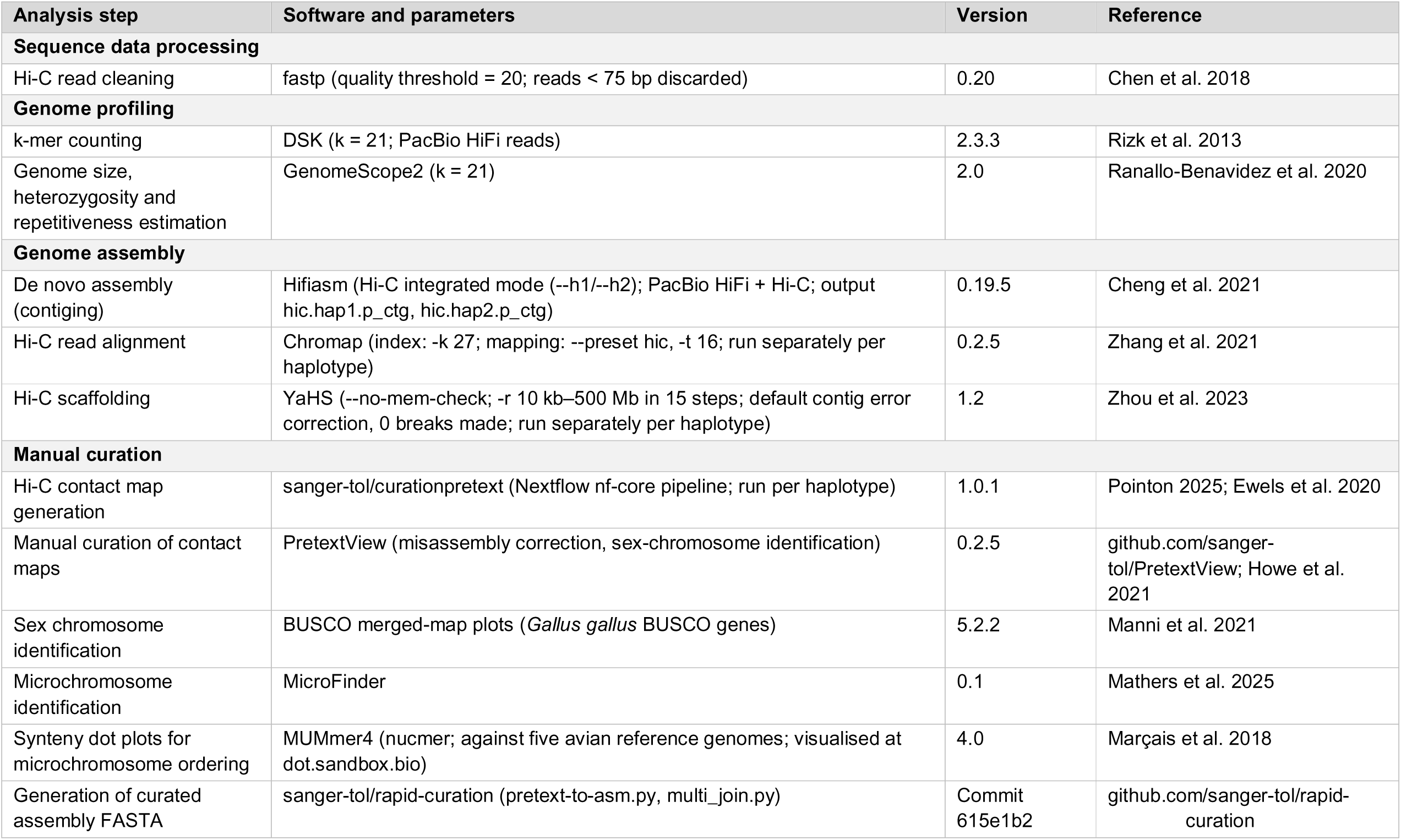
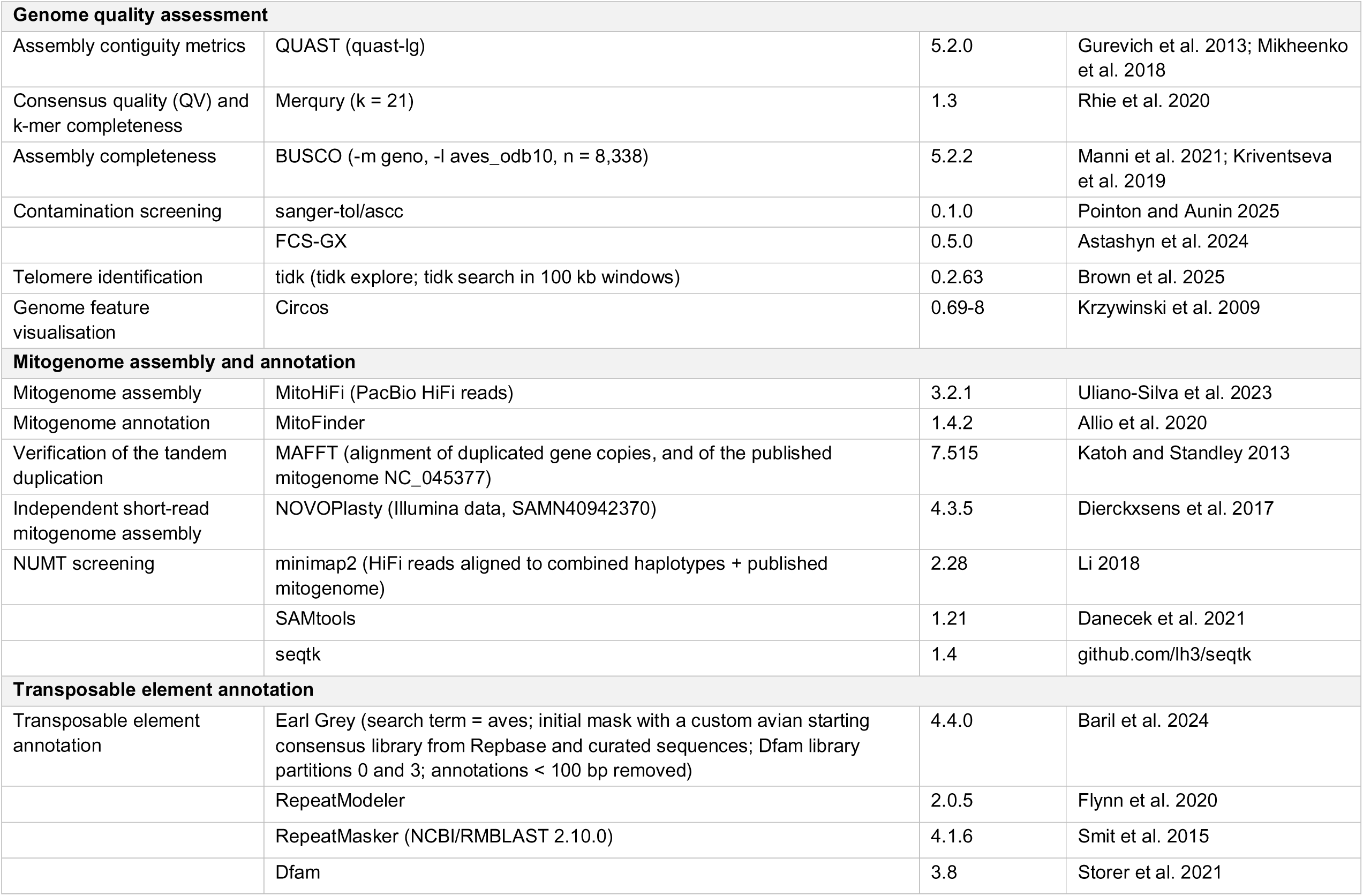
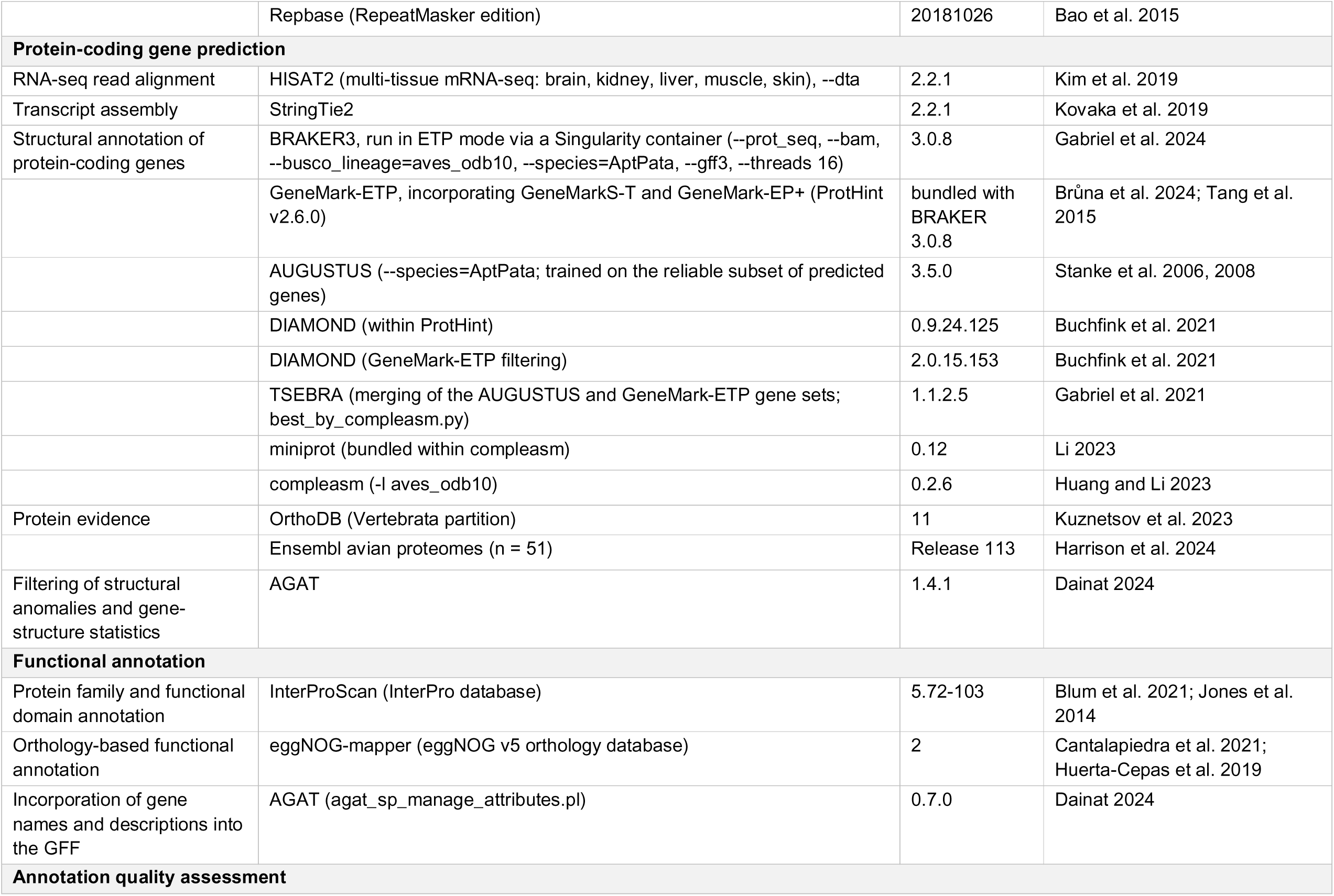
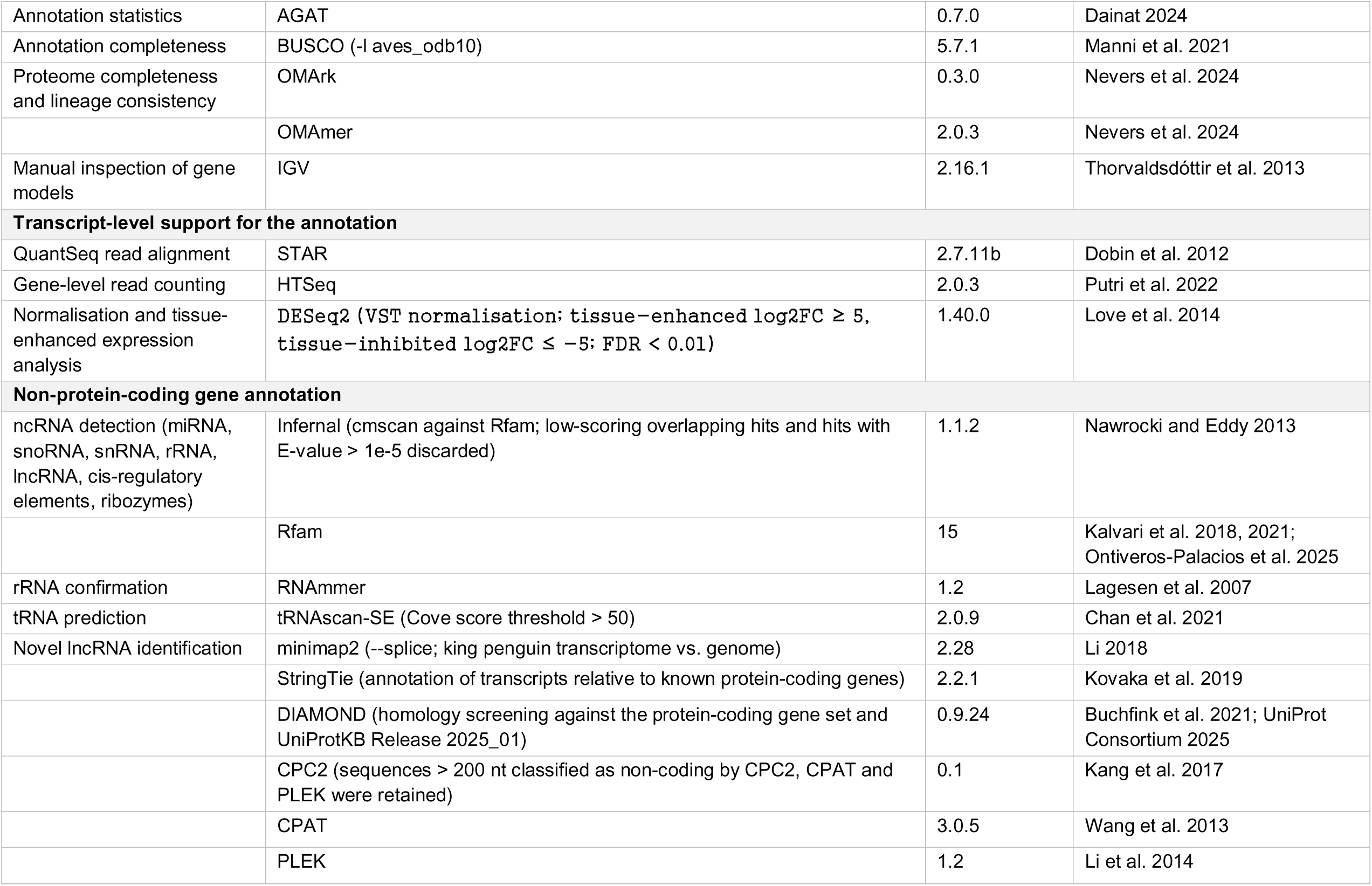
Software used to assemble and annotate the king penguin (*Aptenodytes patagonicus*) genome, together with version numbers and the specific parameters used.

Descriptive statistics and contiguity of the assembly were calculated using quast-lg (Mikheenko et al., 2018) in QUAST (Gurevich et al., 2013). Completeness and quality of the assembly was assessed using Merqury (Rhie et al., 2020) and BUSCO (Manni et al., 2021) against the aves_odb10 dataset (n=8,338) (Kriventseva et al., 2019). Contamination in the assembly was screened with ASCC (github.com/sanger-tol/ascc; Pointon et al., 2025), informed by results from FCS-GX (Astashyn et al., 2024). Telomeric sequences were identified with tidk (Brown et al., 2025): tidk explore was used to identify potential telomeric repeat sequences, and tidk search was run in 100 kb windows. Genome assembly features were plotted using Circos (Krzywinski et al., 2009).

### Mitogenome assembly

We used MitoHiFi (Uliano-Silva et al., 2023) with MitoFinder (Allio et al., 2020) to assemble and annotate the mitogenome from the HiFi reads. The current king penguin mitogenome (NC_045377; Du et al., 2019) is 17,477 bp and contains the standard 37 vertebrate mitochondrial genes. Our assembly produced a circularised mitogenome that was considerably longer (20,520 bp) due to a repetitive tandem duplication (TD) (see Results). The TD was assessed from HiFi read alignments to the assembled mitogenome. Because linearising a circular sequence causes reads crossing the origin to be soft-clipped and therefore under-counts depth towards the ends of the representation, reads were mapped to a doubled reference and coordinates folded back onto the 20,520 bp sequence. Read depth was summarised across the single-copy region and across each copy of the duplicated block, and reads spanning each junction were counted, requiring at least 1 kb of contiguous alignment on both sides. Duplicated gene copies were aligned against each other using MAFFT (Katoh & Standley, 2013). For comparison, we also assembled the mitogenome from Illumina whole-genome sequencing data (SAMN40942370; 118,585,357 read pairs totalling 32.7 Gb, approximately 24× coverage of the 1.35 Gb assembly) using NOVOPlasty v4.3.5 (Dierckxsens et al., 2017), which assembled 5,058 reads into 21,325 bp of contigs, corresponding to approximately 33× mitochondrial coverage.

During contamination screening, we identified scaffold 598 in the bAptPat1.alt assembly as having high mitochondrial similarity. Because the scaffold was approximately twice the length of the published mitogenome (34,801 bp), we assessed whether it represented a mitogenome containing heteroplasmy by aligning it to the published sequence with MAFFT. We additionally screened for potential nuclear-mitochondrial segments (NUMTs) in both haplotypes of the assembly using the approach described in Sozzoni, Ferrer Obiol et al. (2023). Briefly, HiFi reads were aligned with minimap2 (Li, 2018) to a combined reference containing both assembly haplotypes and the published mitogenome, queried using samtools (Danecek et al., 2021) and seqtk (github.com/lh3/seqtk). Reads aligning to both the assembly and mitogenome were realigned to both assembly haplotypes to score putative NUMTs.

### Transposable element annotation

EarlGrey was used to perform transposable element (TE) annotation (Baril et al., 2024), which uses RepeatModeler (Flynn et al., 2020) and RepeatMasker (Smit et al., 2015) with NCBI/RMBLAST. We performed an initial mask using a custom starting consensus library comprising avian sequences derived from Repbase (Bao et al., 2015) and additional curated sequences (Peona et al. 2021; Supplementary Table 2). After the initial mask, EarlGrey was run using library partitions 0 and 3 from Dfam 3.8 (Storer et al., 2021), and Repbase libraries (RepeatMasker edition, version 20181026; Bao et al., 2015), using aves as the search term. Spurious annotations < 100 bp were removed.

### Gene prediction and functional annotation

Structural annotation of protein-coding genes was performed using BRAKER3 (Gabriel et al., 2024; Stanke et al., 2006, 2008). We mapped a multi-tissue (brain, kidney, liver, muscle, and skin) mRNA-seq dataset (Paris et al., 2025; Supplementary Table 3) to the genome using HISAT2 (Kim et al., 2019), and assembled using StringTie2 (Kovaka et al., 2019). BRAKER3 uses GeneMark-ETP (Brůna et al., 2024) to process assembled RNA-seq alignments, which were internally used by GeneMarkS-T (Tang et al., 2015) to screen for potential genes. DIAMOND (Buchfink et al., 2021) searches were incorporated into protein evidence for GeneMark-EP+ to filter the genes, and GeneMark-ETP also performed *ab initio* gene predictions based on self-training. AUGUSTUS (Stanke et al., 2006, 2008) was trained on the reliable subset of predicted genes, and the final gene set derived from AUGUSTUS and GeneMark-ETP was merged using TSEBRA (Gabriel et al., 2021) using miniprot (Li, 2023) and compleasm (Huang & Li, 2023) against the BUSCO aves_odb10 dataset.

We also included protein data: the Vertebrata partition of OrthoDB v11 (Kuznetsov et al., 2023) and 51 avian proteomes from Ensembl Release 113 (Harrison et al., 2024; Supplementary Table 4). AGAT (Dainat, 2024) was used to provide basic filtering for structural anomalies and to quantify statistics regarding structural aspects of the protein-coding regions.

To obtain functional annotations for the predicted protein-coding genes, we searched their sequences against the InterPro protein database using InterProScan (Blum et al., 2021; Jones et al., 2014) and against the eggNOG v5 orthology database (Huerta-Cepas et al., 2019) using eggNOG-mapper (Cantalapiedra et al., 2021). eggNOG gene names and gene descriptions were incorporated into the GFF annotation using the agat_sp_manage_attributes.pl script in AGAT (Dainat, 2024).

Gene predictions and functional annotations were assessed throughout the annotation process using AGAT, BUSCO (Manni et al., 2021), and OMArk with OMAmer (Nevers et al., 2024). Predicted annotations were also manually inspected using IGV (Thorvaldsdóttir et al., 2013).

To assess transcript-level support for the annotation, we analysed a complementary multi-tissue QuantSeq dataset, originating from the same king penguin samples described above (Paris et al., 2025). The dataset was mapped to the genome (Supplementary Table 3) using STAR (Dobin et al., 2013) and gene features were counted using HTSeq (Putri et al., 2022). A PCA was produced following a VST normalisation on the counts table to assess tissue-specific expression patterns. To count the number of protein-coding genes supported by the QuantSeq data in each tissue (≥5 replicates per tissue), we quantified the normalised gene expression in bins of 0, 0-50, 50-100, 100-200, 200-500, ≥500. We then performed an analysis of tissue-enhanced gene expression as per Paris et al., (2025) with DESeq2 (Love et al., 2014), using a log2foldchange ≥ 5 to determine significant tissue-enhanced transcripts (significantly higher in a particular tissue compared to the average level in all other tissues) and ≤ −5 as tissue-inhibited transcripts (significantly lower in a particular tissue compared to the average level in all other tissues). An FDR threshold < 0.01 was used for significance.

### Non-protein-coding gene annotation

miRNAs, snoRNAs, snRNAs, rRNA and databased long-non-coding RNAs (lncRNAs) were detected using a combination of Infernal (Nawrocki & Eddy, 2013) and Rfam (Kalvari et al., 2018, 2021) by searching the Rfam 15 database (Ontiveros-Palacios et al., 2025) with cmscan. Low-scoring overlapping hits and hits with an E-value > 1e-5 were discarded, and the remaining ncRNAs were grouped into non-protein-coding RNA categories. We additionally annotated cis-regulatory elements (CREs) and Ribozymes using the Rfam database. rRNAs and their subunits were confirmed using RNAmmer (Lagesen et al., 2007). tRNAs were predicted using tRNAScan-SE (Chan et al., 2021). To compare our results to other avian species, we followed the approaches of Ottenburghs et al., (2021) by applying a Cove score threshold value of 50 to remove low-quality tRNA gene predictions. To further assess the quality of the tRNA annotation, we additionally analysed the tRNA content of *Spheniscus humboldti* (GCA_027474245.1).

Novel lncRNAs were identified by aligning the king penguin transcriptome (Paris et al., 2025) to the genome using minimap2 --splice (Li, 2018). Resulting alignments were used as input to StringTie (Kovaka et al., 2019) to annotate transcripts relative to known protein-coding genes, designating unannotated alignments as potential novel transcripts. To further exclude potential coding sequences, we used DIAMOND (Buchfink et al., 2021) to screen novel transcripts for homology against the protein-coding gene set and the UniProtKB Release 2025_01 (Swiss-Prot & TrEMBL; UniProt Consortium, 2025). To distinguish lncRNAs from protein-coding RNAs, we applied CPC2 (Kang et al., 2017), CPAT (Wang et al., 2013), and PLEK (Li et al., 2014), retaining only those sequences >200 nt in length and classified as non-coding by all three programs.

### DNA methylation

Average DNA methylation was calculated in 50 kb non-overlapping windows, based on the mean methylation at validated genome-wide CpG loci measured in 64 individuals (Cristofari et al., 2026). Briefly, DNA extracted from whole blood was converted using the EMseq enzymatic conversion method (Vaisvila et al., 2021) and sequenced on a NovaSeq 6000. After read preprocessing, we used bsbolt (Farrell et al., 2021) to align the reads to the genome, and to call methylation levels. See Cristofari et al., (2026) for further methodological details.

## Results

### Genome assembly

*K*-mer profiling (*k* = 21) generated an estimated genome span of 1.15 Gb, with a heterozygosity of 0.6% (Supplementary Figure 1). The preliminary scaffolded assembly produced two phased haplotypes comprising 685 scaffolds in haplotype 1 and 733 in haplotype 2. Initial inspection indicated a highly contiguous genome, including clear visual identification of the sex chromosomes (Supplementary Figure 2). After manual curation, the final assembled genome comprised two phased haplotypes (bAptPat1.pri and bAptPat1.alt), including 32 autosomes, plus the Z and W chromosomes. A previous karyotype study reported that the species possesses 36 chromosomes, including eight macrochromosomes and 28 microchromosomes (Cardozo et al., 2003). Defining microchromosomes as <35 Mb and dot chromosomes as <5 Mb (Huang et al., 2023; Waters et al., 2021), we recovered nine macrochromosomes (including the Z and W sex chromosomes), 18 microchromosomes, and seven dot chromosomes. Our inability to fully resolve the karyotype was due to the highly fragmented nature of the dot chromosomes, whose associated scaffolds were too small to be reliably visualised using Hi-C. Autosomal macrochromosomes ranged in length from 74 Mb to 227 Mb, microchromosomes ranged in length from 6.48 Mb to 25.76 Mb, and dot chromosomes ranged in length from 0.34 Mb to 3.2 Mb. We identified telomeric regions on at least one end of 16 of the 34 assembled chromosomes (Supplementary Figure 3). We successfully assembled the Z (90 Mb) and W chromosomes (48 Mb), including visual identification of the pseudoautosomal region (PAR; Supplementary Figure 2), providing a promising resource to further investigate gene content, transposable element activity, the accumulation of mutational load in bird sex chromosomes (e.g. Peona et al., 2021; Xu et al., 2019), and the evolution of potential neo-sex chromosomes in penguins (Gunski et al., 2017).

The final assembly spans 1.35 Gb, with 95% of the bases captured in the 34 assembled chromosomes (Figure 1). The genome is high quality: assessed across both haplotypes combined, the assembly has a per-base consensus quality value (QV) of 63.8 (0.414 base errors per Mb) and a *k*-mer completeness of 99.68%. The primary haplotype (bAptPat1.pri) has a BUSCO completeness of 97.2% (aves_odb10). Per-haplotype values for all metrics are given in Table 2. Compared to the previous draft-level assembly (BGI_Apat.V1), the presented chromosome assembly (bAptPat1.pri) increased the scaffold N50 by ∼30-fold (from 2.9 Mb to 86.6 Mb) and reduced the scaffold count by 26-fold (from 15,968 to 618). The current assembly exceeds the standards outlined by both the Earth BioGenome Project (Lewin et al., 2018), and the Vertebrate Genomes Project (Rhie et al., 2021).

**Figure 1.**
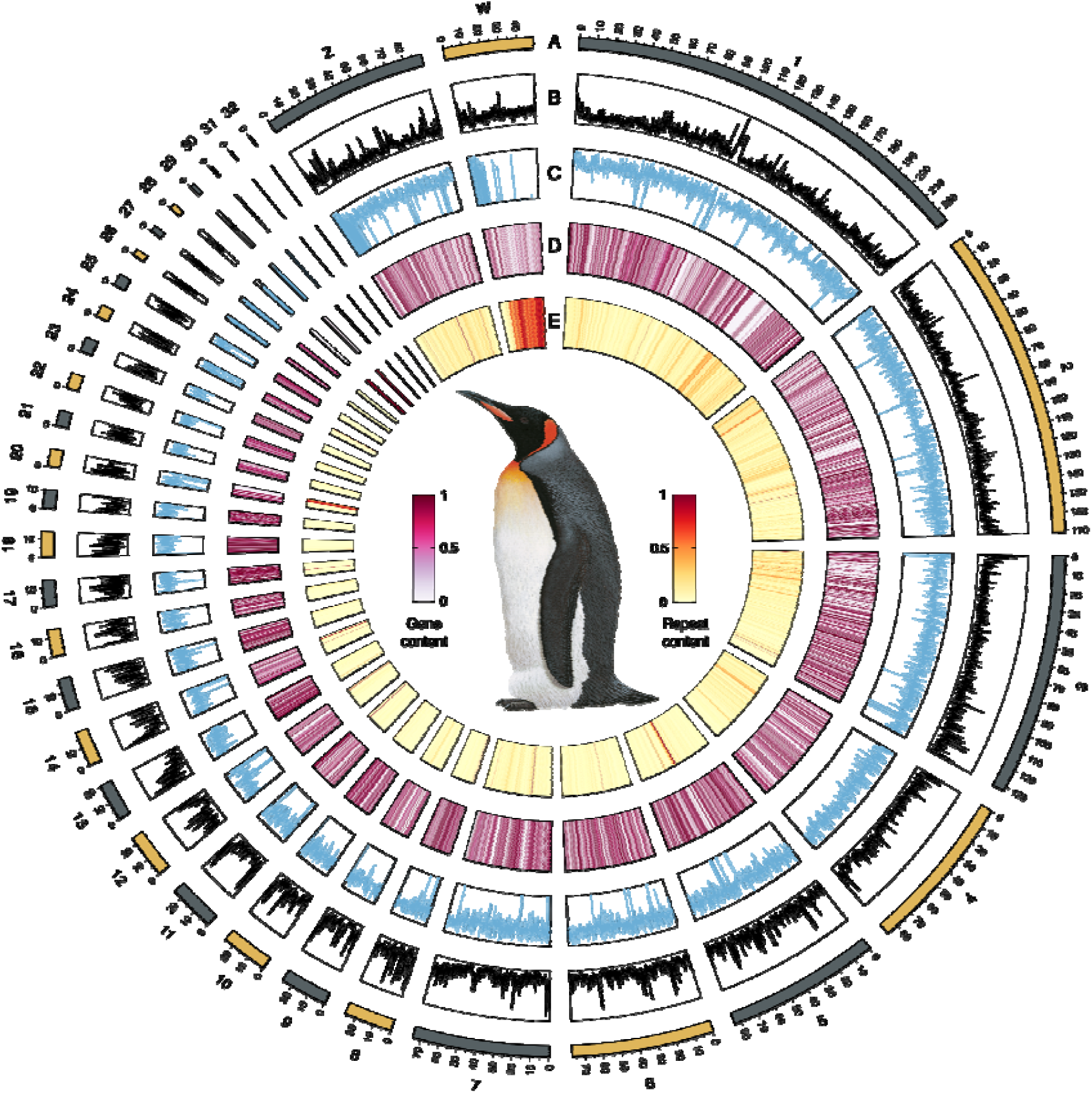
Genomic features of the king penguin (*Aptenodytes patagonicus*) assembly ‘Pen/Se-guin’. Circos plot of genome characteristics, showing from the outside to the inside: (**A**) chromosome ideograms; (**B**) GC content in 50 kb windows; (**C**) methylation marks in 50 kb windows; (**D**) protein-coding gene content in 50 kb windows; and (**E**) repeat content in 50 kb windows. The image of the king penguin is reproduced with permission from © Lynx Nature Books.

**Table 2.**
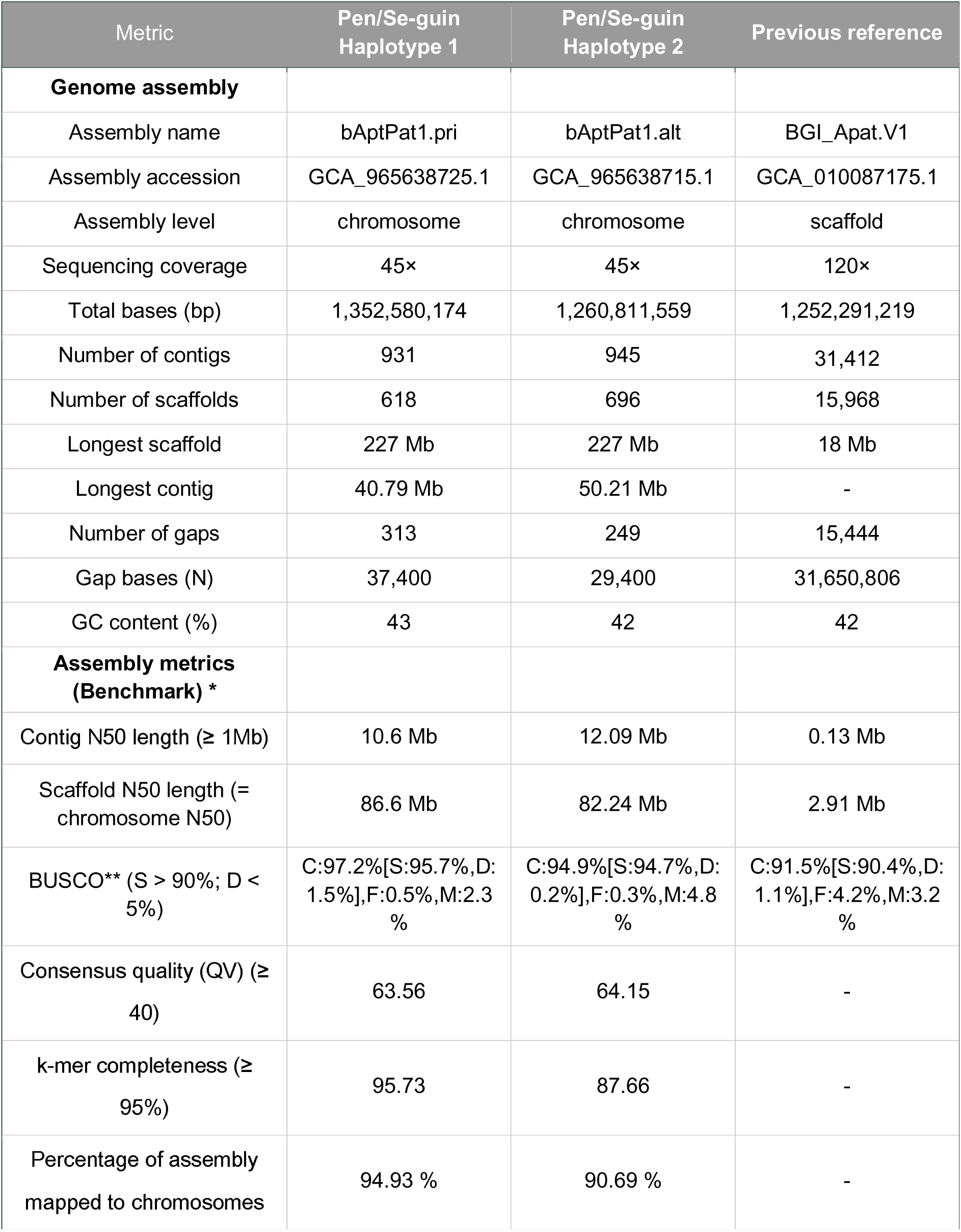
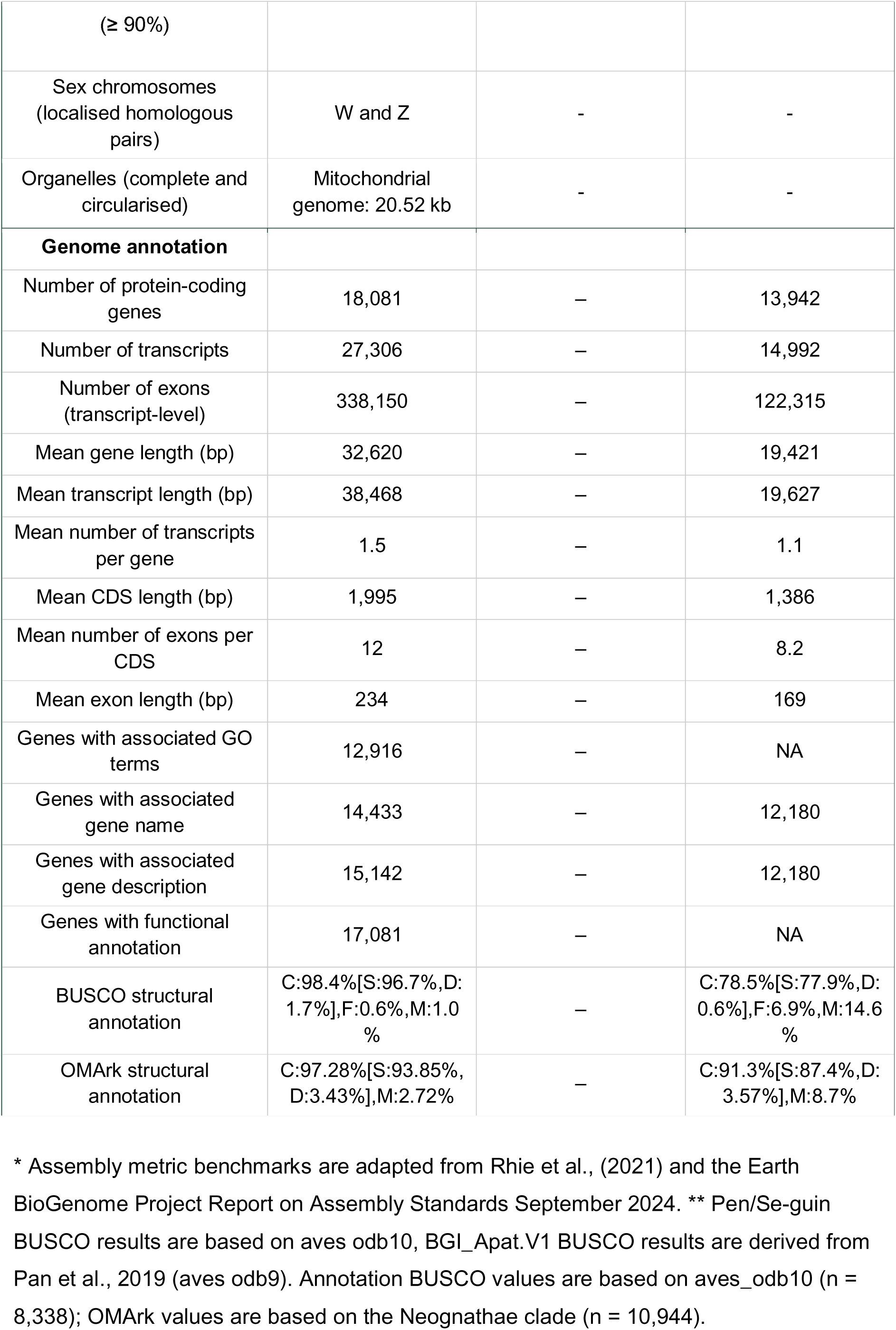
Genome assembly and annotation statistics for the king penguin (*Aptenodytes patagonicus*). Statistics are given for the presented assembly - Pen/Se-guin - and compared to the previous reference genome (BGI_Apat.V1). Haplotype 1 statistics are for the nuclear assembly and exclude the circularised mitochondrial genome (20,520 bp), which is reported separately below; they therefore correspond directly to the NCBI record for GCA_965638725.1. Gene annotation was performed on haplotype 1 (bAptPat1.pri) only.

| Metric | Pen/Se-guin<br>Haplotype 1 | Pen/Se-guin<br>Haplotype 2 | Previous reference |
| --- | --- | --- | --- |
| **Genome assembly** |  |  |  |
| Assembly name | bAptPat1.pri | bAptPat1.alt | BGI_Apat.V1 |
| Assembly accession | GCA_965638725.1 | GCA_965638715.1 | GCA_010087175.1 |
| Assembly level | chromosome | chromosome | scaffold |
| Sequencing coverage | 45× | 45× | 120× |
| Total bases (bp) | 1,352,580,174 | 1,260,811,559 | 1,252,291,219 |
| Number of contigs | 931 | 945 | 31,412 |
| Number of scaffolds | 618 | 696 | 15,968 |
| Longest scaffold | 227 Mb | 227 Mb | 18 Mb |
| Longest contig | 40.79 Mb | 50.21 Mb | - |
| Number of gaps | 313 | 249 | 15,444 |
| Gap bases (N) | 37,400 | 29,400 | 31,650,806 |
| GC content (%) | 43 | 42 | 42 |
| **Assembly metrics**<br>**(Benchmark) \*** |  |  |  |
| Contig N50 length (≥ 1Mb) | 10.6 Mb | 12.09 Mb | 0.13 Mb |
| Scaffold N50 length (= chromosome N50) | 86.6 Mb | 82.24 Mb | 2.91 Mb |
| BUSCO** (S > 90%; D < 5%) | C:97.2%[S:95.7%,D:1.5%],F:0.5%,M:2.3% | C:94.9%[S:94.7%,D:0.2%],F:0.3%,M:4.8% | C:91.5%[S:90.4%,D:1.1%],F:4.2%,M:3.2% |
| Consensus quality (QV) (≥ 40) | 63.56 | 64.15 | - |
| k-mer completeness (≥ 95%) | 95.73 | 87.66 | - |
| Percentage of assembly mapped to chromosomes | 94.93 % | 90.69 % | - |
| (≥ 90%) |  |  |  |
| Sex chromosomes<br>(localised homologous<br>pairs) | W and Z | - | - |
| Organelles (complete and<br>circularised) | Mitochondrial<br>genome: 20.52 kb | - | - |
| **Genome annotation** |  |  |  |
| Number of protein-coding<br>genes | 18,081 | – | 13,942 |
| Number of transcripts | 27,306 | – | 14,992 |
| Number of exons<br>(transcript-level) | 338,150 | – | 122,315 |
| Mean gene length (bp) | 32,620 | – | 19,421 |
| Mean transcript length (bp) | 38,468 | – | 19,627 |
| Mean number of transcripts<br>per gene | 1.5 | – | 1.1 |
| Mean CDS length (bp) | 1,995 | – | 1,386 |
| Mean number of exons per<br>CDS | 12 | – | 8.2 |
| Mean exon length (bp) | 234 | – | 169 |
| Genes with associated GO<br>terms | 12,916 | – | NA |
| Genes with associated<br>gene name | 14,433 | – | 12,180 |
| Genes with associated<br>gene description | 15,142 | – | 12,180 |
| Genes with functional<br>annotation | 17,081 | – | NA |
| BUSCO structural<br>annotation | C:98.4%[S:96.7%,D:<br>1.7%],F:0.6%,M:1.0<br>% | – | C:78.5%[S:77.9%,D:<br>0.6%],F:6.9%,M:14.6<br>% |
| OMArk structural<br>annotation | C:97.28%[S:93.85%,<br>D:3.43%],M:2.72% | – | C:91.3%[S:87.4%,D:<br>3.57%],M:8.7% |

\* Assembly metric benchmarks are adapted from Rhie et al., (2021) and the Earth BioGenome Project Report on Assembly Standards September 2024. \*\* Pen/Se-guin BUSCO results are based on aves odb10, BGI\_Apat.V1 BUSCO results are derived from Pan et al., 2019 (aves odb9). Annotation BUSCO values are based on aves\_odb10 (n = 8,338); OMArk values are based on the Neognathae clade (n = 10,944).

### Mitogenome assembly and annotation

The assembled circularised mitogenome is 20,520 bp in length and includes the full set of 37 vertebrate mitochondrial genes, a tandem duplication containing three tRNA genes (tRNA-Thr, tRNA-Pro and tRNA-Glu), a degenerate copy of ND6, and a second copy of the control region (Supplementary Table 5). The duplicated tRNA genes showed 100% sequence similarity, whereas the first ND6 copy is degenerate and was annotated as a pseudogene. We note that the duplicated ND6 was not identified in the automated annotation performed using MitoFinder and was manually curated using sequence alignments. The tandem duplication was supported directly by the HiFi read alignments (Supplementary Figure 4): 20 reads spanned the junction between CYTB and the first copy of the duplicated block, and 21 spanned the junction between the two copies, each with at least 1 kb of contiguous alignment on either side, while 14 reads traversed the entire duplicated block in a single contiguous alignment. Mean read depth was 28.4× across the single-copy region, and 21.3× and 29.3× across copies 1 and 2 of the duplicated block, respectively. Circularity was supported by HiFi reads spanning the point of circularisation (29 of 46 mitochondrial reads; Supplementary Figure 4).

The observed gene order is in agreement with the putative ancestral avian pattern (Montaña-Lozano et al., 2022), which typically contains a pseudogene Ψ (a degenerate copy of CYTB - the CYTB pseudogene was not annotated here), four functional genes (T, P, ND6 and E), and an extended control region (Urantówka et al., 2020). The mitochondrial gene order is consistent with the arrangement reported in Procellariiformes, the evolutionarily closest order to Sphenisciformes (Abbott et al., 2005; Morris-Pocock et al., 2010). The final mitogenome is 3,043 bp longer than the previously published mitogenome generated using long-range PCR (17,477 bp). Despite mitochondrial coverage (∼33×) comparable to that of the HiFi data (∼28×), the short-read assembly (NOVOplasty) did not circularise and poorly resolved the duplicated region, reflecting the inability of short reads to span the ∼2.4 kb near-identical repeat rather than insufficient sequencing depth. By aligning the mitogenome assemblies (MitoHiFi, NC_045377, NOVOplasty), we found that the initially identified duplication (see Materials and Methods) was absent in the published mitogenome and was poorly assembled in the short-read assembled mitogenome. It has been previously reported that more complex mitogenomes are more problematic to assemble, and the observation of more contiguous mitogenomes recovered from long-read sequencing technologies has been attributed to repetitive duplications (Formenti et al., 2021). This fully characterised mitogenome offers a resource to further explore the potential for selection on penguin mitochondrial genes (Ramos et al. 2018).

Finally, we identified putative NUMTs on seven chromosomes and found a particular enrichment of NUMTs on the W chromosome and on scaffold 598 in haplotype 2 (bAptPat1.alt) (Supplementary Table 6). Together, this analysis highlights that mitogenomes generated from long-read data are more complete than many published mitogenomes based on short-read or long-range PCR and Sanger sequencing methods (Formenti et al., 2021) and that avian, and other assembled mitogenomes, need to be carefully confirmed and curated to avoid the accessioning of incomplete or inaccurate mitogenomes (Sangster & Luksenburg, 2021).

### Transposable element annotation

TE annotation produced a curated library of 238 consensus sequences. Overall, 16.3% of the genome was characterised as repetitive (Supplementary Table 7), with LINEs comprising the most abundant transposable element class (5.6%), consistent with patterns observed in avian genomes (Peona et al., 2022).

### Protein-coding genes

Protein-coding gene prediction with BRAKER3 generated 18,081 predicted gene models, with a mono-to-multi-exon gene ratio of 0.162, a BUSCO completeness of 98.4%, and an OMArk completeness of 97.28% (see Table 2 for full annotation statistics). Within OMArk, the proteome showed an 85% consistent lineage placement. No contamination was identified. Functional annotation showed that 94.47% of the protein-coding genes have an annotation (protein family and functional domain annotation), 83.87% have an associated gene description, and 79.84% have an associated gene name. Overall, the protein-coding annotation represents a substantial improvement over the current annotation for the species (Pan et al., 2019).

Using the QuantSeq RNA dataset, we found that 83% of predicted protein-coding genes were supported by at least one RNA-sequencing read. PCA of tissue-level expression profiles was highly consistent with previous analyses based on the king penguin *de novo* transcriptome (Figure 2A; Paris et al., 2025), indicating that the annotation captures biologically meaningful patterns of tissue-specific expression. Across tissues, 67–77% of genes showed detectable expression (>0) in at least half of the replicates (5 of 10), while 8–11% exhibited high expression levels (>500; Figure 2B). Tissue-enhanced and tissue-inhibited genes were identified across all macro- and microchromosomes, including several putative tissue-specific gene clusters (Figure 2C). Together, these results support the high quality and biological relevance of the king penguin genome annotation and provide a foundation for future intra- and inter-specific transcriptomic studies (Paris et al., 2025).

**Figure 2.**
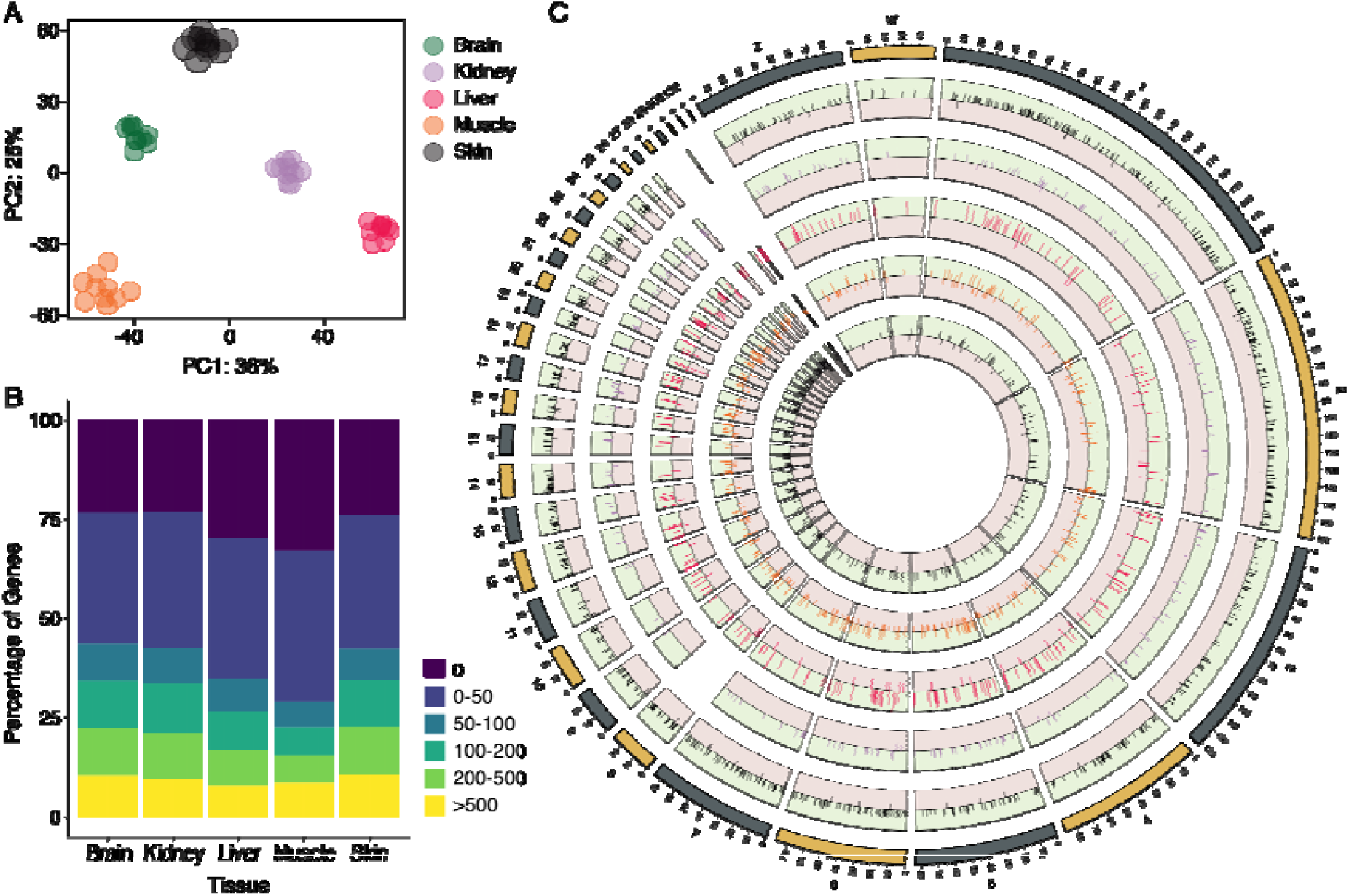
Characteristics of the protein-coding gene annotation for the king penguin (*Aptenodytes patagonicus*) assembly ‘Pen/Se-guin’ (bAptPat1.hap1.1). (**A**) Principal component analysis (PCA) of variance-stabilising transformation (VST) gene expression counts for five tissues: brain (green), kidney (purple), liver (pink), muscle (orange), and skin (black). (**B**) Stacked bar plot showing the percentage of genes in six VST expression bins (0, 0–50, 50–100, 100–200, 200–500, and >500) across the five tissues. (**C**) Circos plot showing tissue-enhanced and tissue-suppressed genes across the five tissues. See Table 2 for further information on the protein-coding gene annotation.

### Non-protein-coding gene annotation

Overall, we annotated six types of non-coding RNAs (Supplementary Table 8), including 281 miRNAs, 253 snoRNAs, 60 snRNAs, 341 rRNAs (Supplementary Table 9), 358 tRNAs (Supplementary Table 10), and 16 known lncRNAs. Using the king penguin transcriptome aligned to the genome, we identified an additional 164 lncRNAs, with an average length of 673 bp. lncRNAs were located throughout the genome but were particularly enriched on the macrochromosomes and the Z chromosome. We additionally predicted 81 CREs and six ribozymes. tRNA detection is highly dependent on genome assembly quality, and we identified more than double the number of tRNAs in comparison to *Pygoscelis adeliae* (n = 166) and *Aptenodytes forsteri* (n = 135) (Ottenburghs et al., 2021). The scaffold-level genomes of *P. adeliae* and *A. forsteri* were both assembled using short-read data and show lower scaffold N50s: 5.07 Mb and 5.11 Mb, respectively. To evidence if the higher number of identified tRNAs in the king penguin genome may be due to increased genome assembly contiguity, we also analysed the tRNA content of the *Spheniscus humboldti* chromosome-level assembly (GCA_027474245.1), finding 389 tRNAs (Cove threshold > 50). This suggests that improved genome assembly quality improves the ability to annotate tRNA genes.

## Discussion

Compared to the previous draft-level assembly of the king penguin, the assembly presented here (‘Pen/Se-guin’) increases contiguity ∼30-fold, is haplotype-resolved at chromosome level, and is accompanied with a substantially improved annotation. It is also only the third chromosome-level genome available for any penguin, and the first for the genus *Aptenodytes*, extending the taxonomic breadth available for comparative analyses within an iconic order of birds with highly specialised adaptations to deep-ocean ecology and considerable conservation concern (Boersma, 2008; Barbraud et al., 2020; Ropert-Coudert et al., 2019; Cole et al., 2022).

Decades of individual-based monitoring in the king penguin have generated longitudinal records of survival, breeding phenology, reproductive success and foraging behaviour, alongside extensive behavioural, demographic and physiological data (Le Bohec et al., 2007; Bardon et al., 2023, 2026), which can now be interrogated at much higher genomic resolution. Comparable data exist for only a handful of wild vertebrate populations, where combining them with genome-wide data has revealed the genetic basis of inbreeding depression and contemporary selection (e.g. Huisman et al., 2016; Stoffel et al., 2021; Bosse et al., 2017).

Several aspects of king penguin biology are particularly useful for linking genomic variation to fitness. The species exhibits one of the longest breeding cycles of any bird, extending over 14 months (Descamps et al., 2002), resulting in an early–late breeding system in which individuals breed under markedly different environmental conditions (Olsson, 1996; Le Bohec et al., 2007), with late breeders experiencing reduced chick survival (Stier et al., 2014). Adults alternate prolonged fasting periods on land with extended foraging trips at sea (Cherel et al., 1994; Saraux et al., 2012), and foraging efficiency and breeding success are further associated with age, experience, and individual quality (Le Vaillant et al., 2012, 2016). These contrasts provide a natural framework for investigating the genomic and transcriptomic basis of variation in fitness and life-history strategies in the wild (e.g. Nitta Fernandes et al., 2025).

King penguins are also long-lived (average of 20 years in the wild), providing a valuable vertebrate system for investigating ageing and senescence (Cristofari et al., 2026). The assembly has already been used to build an epigenetic clock for the king penguin, which has also been extended as a baseline for several other avian species (Hukkanen et al., 2026). Epigenetic clocks are increasingly used to estimate age and ageing rate in wild vertebrates (Jarman et al., 2015; Heydenrych et al., 2021), and a contiguous, well-annotated reference allows informative CpG sites to be placed in genomic context and linked to gene models. The annotation presented here, combined with known-age individuals from the long-term monitoring programme, makes such analyses feasible in a wild population.

The assembly will also support research on understanding responses to environmental change. King penguin foraging ecology is closely linked to the Antarctic Polar Front (APF) (Bost et al., 1997, 2004; Bost, et al., 2015; Cherel et al., 2018; Scheffer et al., 2010), and climate-driven shifts in the APF are predicted to increase foraging distances, reduce breeding success, and lead to major population declines and/or colony relocations during the coming century (Böning et al., 2008; Le Bohec et al., 2008; Cristofari et al., 2018). Whether populations can respond adaptively, and not only demographically, requires genome-wide data. King penguins possess a large effective population size and exhibit little population structure (Trucchi et al., 2014; Cristofari et al., 2018), demographic characteristics that increase the efficacy of natural selection (Trucchi et al., 2025) and make the species well suited to detecting signatures of contemporary selection. Emerging selective pressures, including the recent spread of highly pathogenic avian influenza (H5N1) into subantarctic and Antarctic regions (Clessin et al., 2025), further highlight the potential for understanding the genomic basis of adaptation and resilience to disease.

Together, this assembly and annotation establish the genomic foundation for using the king penguin to investigate the basis of fitness, ageing and adaptation in a wild, individually monitored vertebrate, while contributing a divergent lineage to comparative studies of penguin responses to environmental change in the Southern Ocean.

## Acknowledgements

The study was approved by the French ethics committee (last: APAFIS#29338-2020070210516365) and the French Polar Environmental Committee and permits to handle animals and access breeding sites were delivered by the “Terres Australes et Antarctiques Françaises” (TAAF). Bioinformatic analyses were performed on the HPC clusters at the Department of Life and Environmental Sciences (“HappyComputing@DiSVA”), Marche Polytechnic University, Italy and at the Wellcome Sanger Institute, UK. We thank Valentina Peona for kindly providing access to curated avian transposable element libraries.

## Data Availability

All analysis scripts and the annotation files are available on GitHub (https://github.com/josieparis/King-penguin-genome-assembly) and archived on Zenodo (https://doi.org/10.5281/zenodo.22968008; Paris et al. 2026).

The genome assembly for the primary haplotype (bAptPat1.pri) can be accessed via the NCBI using the accession number GCA_965638725.1. The alternate haplotype (bAptPat1.alt) can be accessed via the NCBI using the accession number GCA_965638715.1. The NCBI RefSeq assembly is available under accession GCF_965638725.1. The NCBI RefSeq assembly differs from the submitted assembly by the removal of the mitochondrial chromosome.

Sequencing data used for genome assembly are available through the European Nucleotide Archive (ENA) under BioProject accession PRJEB86486 and BioSample accessions SAMEA117789397 (haplotype 1 - primary) and SAMEA117789398 (haplotype 2 - alternate). RNA-seq data are available under the BioProject accession PRJEB64484. DNA methylation data are available under the BioProject accession PRJNA1187342.

The annotated mitochondrial genome can be accessed on NCBI Genbank via the accession number PZ518787.1.

Two annotations are available for this assembly. The annotation described in this manuscript was generated with BRAKER3 using species-specific multi-tissue RNA-seq evidence and is available on the project’s GitHub. Independently, NCBI generated an annotation using the Eukaryotic Genome Annotation Pipeline (Annotation Release GCF_965638725.1-RS_2025_08). This annotation was produced automatically and is maintained and periodically updated by NCBI. All annotation statistics reported here refer to the BRAKER3 annotation.

## Supplementary Figures

**Supplementary Figure 1.**
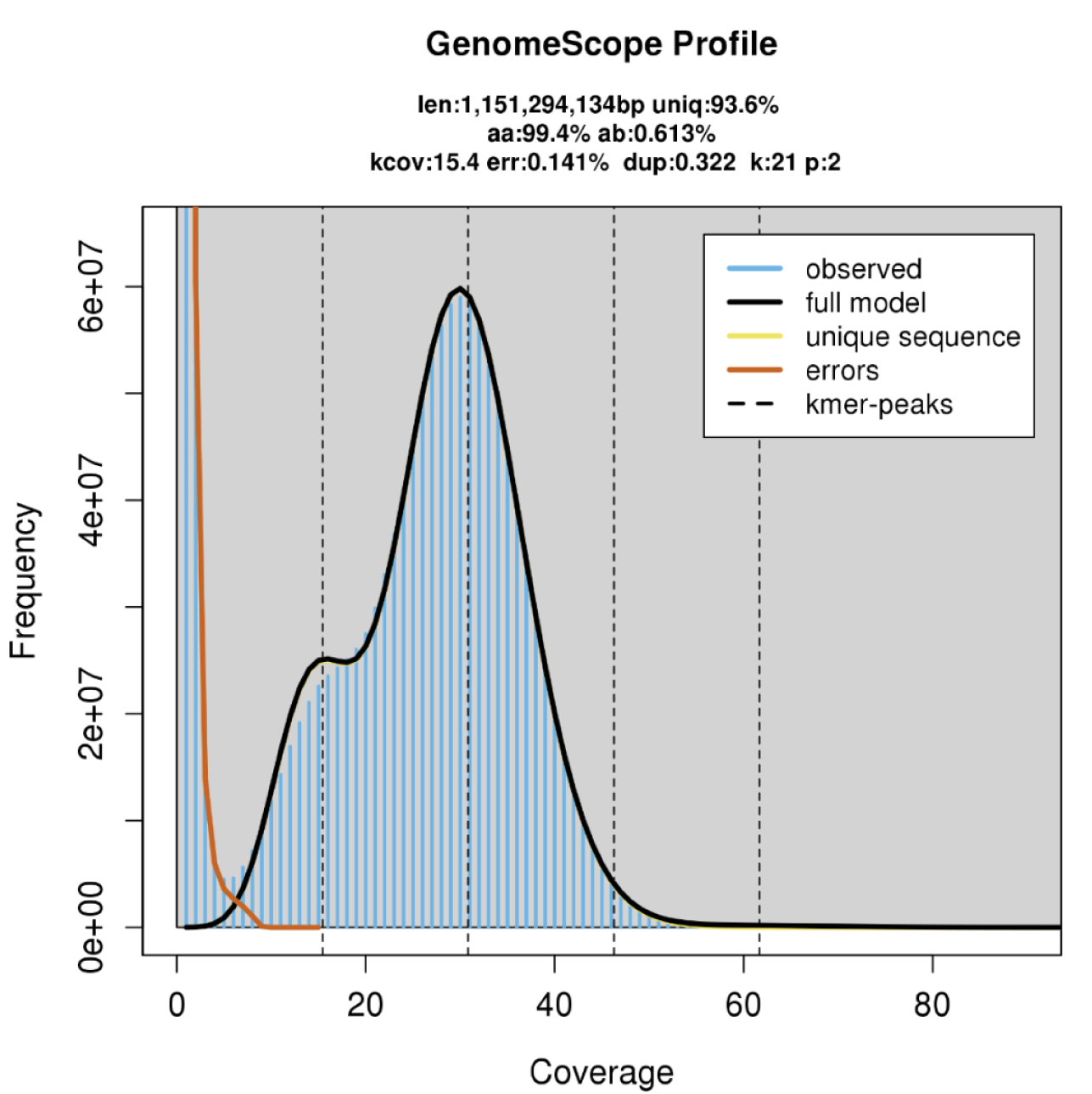
GenomeScope profile of the King penguin (*Aptenodytes patagonicus*) PacBio HiFi reads, computed using 21-mers.

**Supplementary Figure 2.**
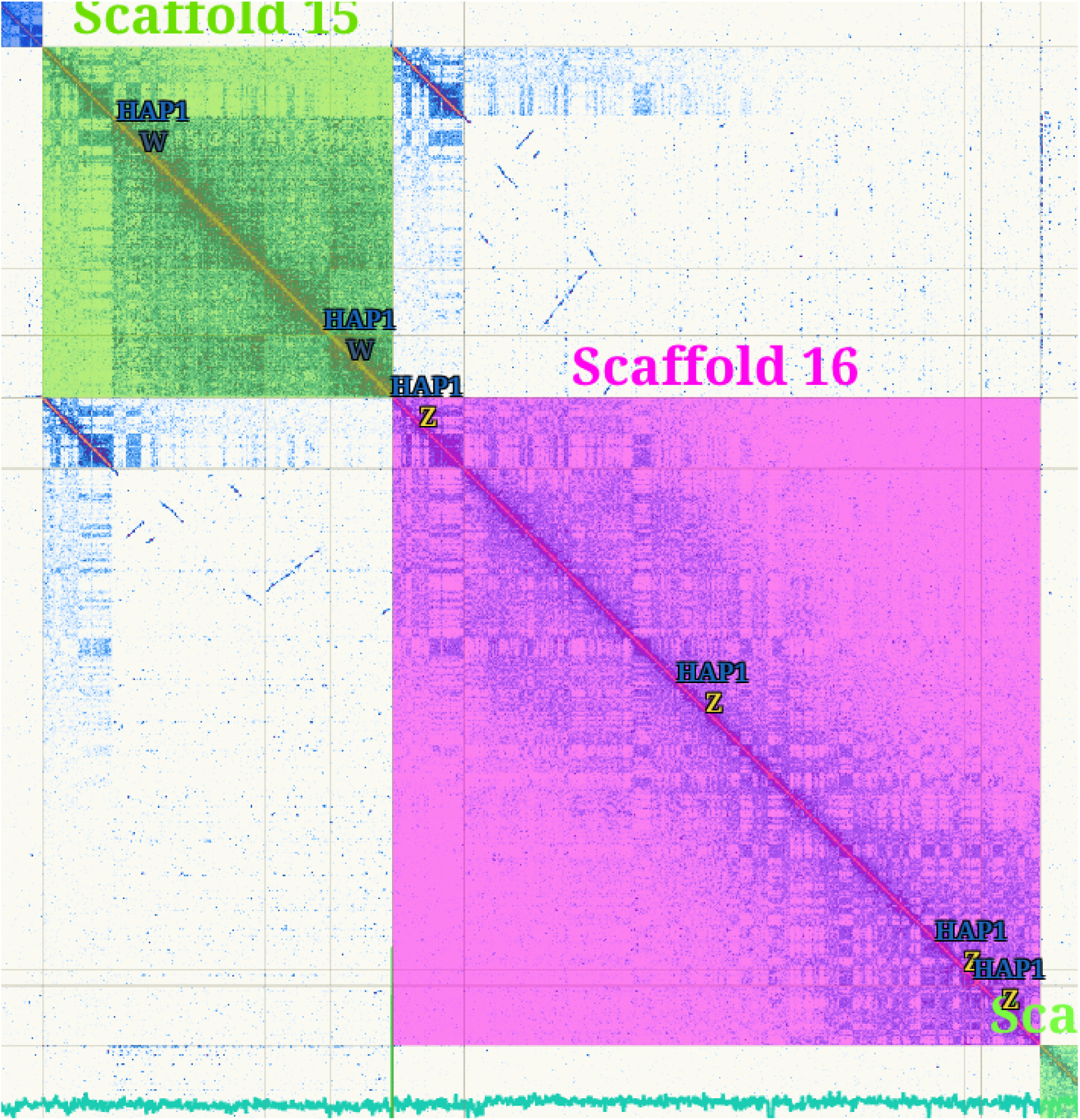
PretextMap showing visual identification of the W and Z chromosomes, including the pseudoautosomal region (PAR) in the king penguin (*Aptenodytes patagonicus*) genome.

**Supplementary Figure 3.**
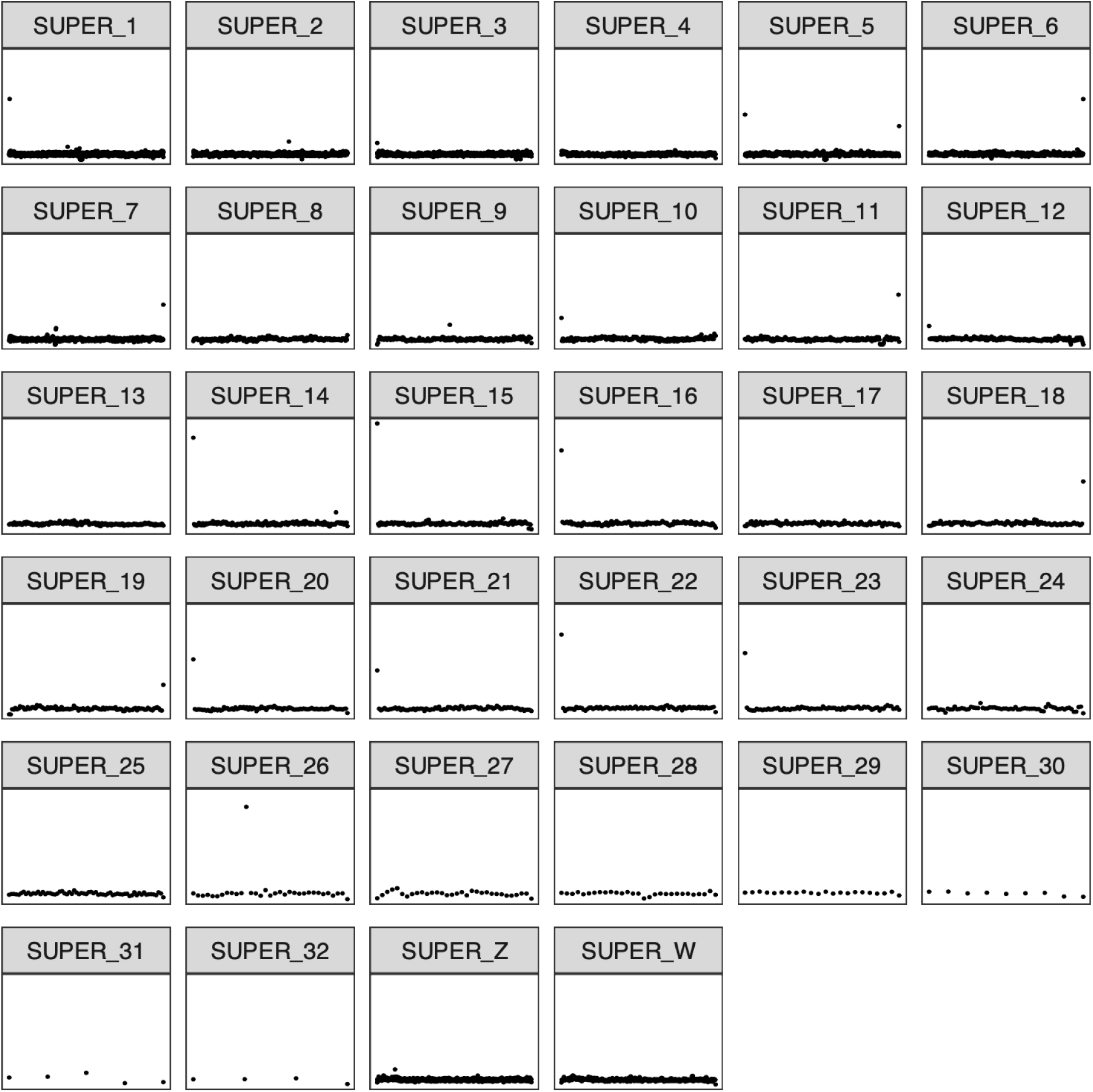
Identification of telomeric repeats (identified with tidk) across the chromosomes of the king penguin (*Aptenodytes patagonicus*) genome.

**Supplementary Figure 4.**
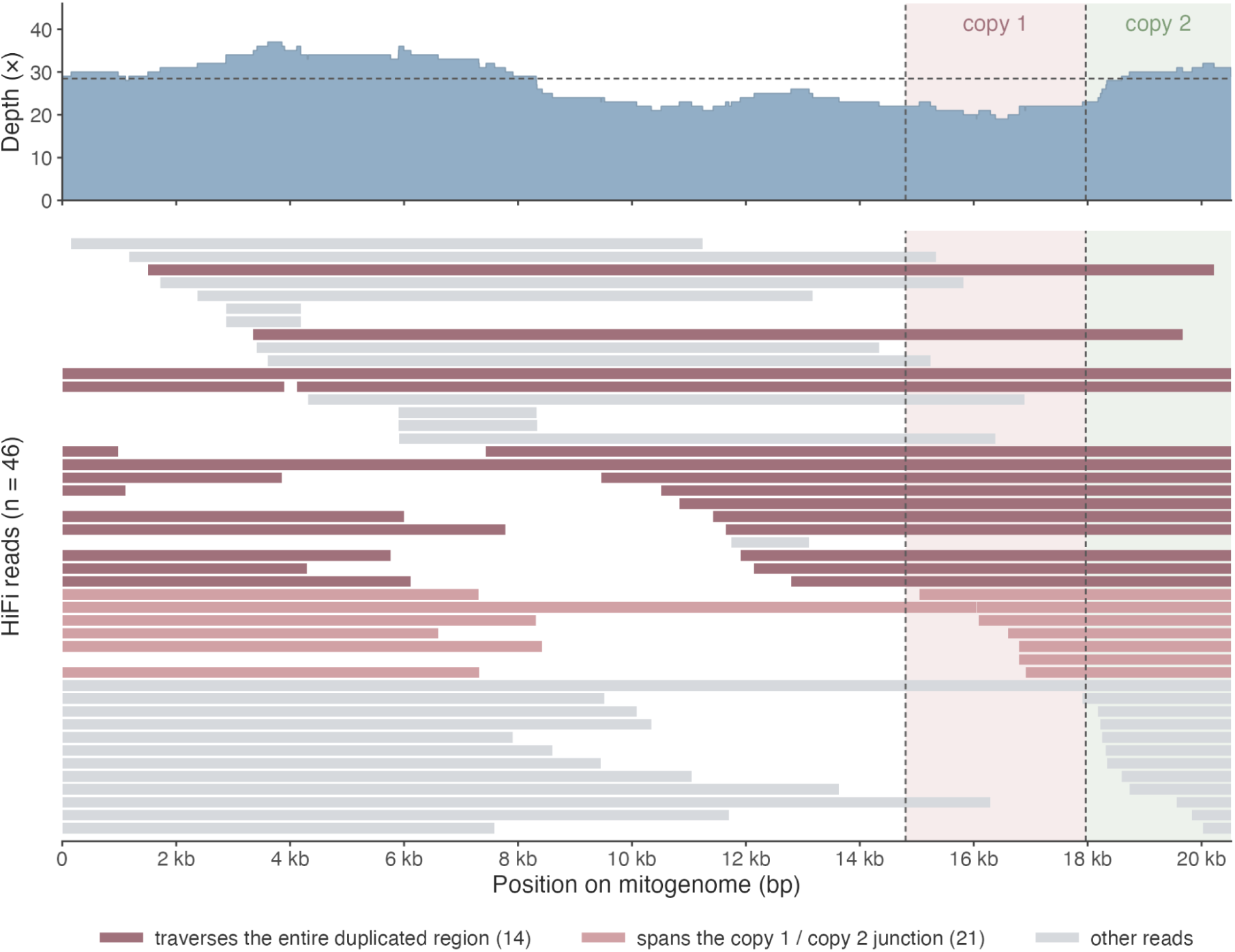
HiFi support for the tandem duplication in the mitogenome. (Top) Per-base HiFi read depth across the 20,520 bp mitogenome. The dashed line marks the mean depth of the single-copy region (positions 1–14,804; 28.4X). (Bottom) Individual HiFi read alignments (n = 46), ordered by start position. Reads spanning the junction between the two copies of the duplicated block (positions 17,967/17,968) with at least 1 kb of contiguous alignment on either side are shown in light red (n = 21). Reads traversing the entire duplicated region in a single contiguous alignment are shown in dark red (n = 14). All other reads are grey. Shading marks copy 1 (14,805-17,967) and copy 2 (17,968-20,520) of the duplicated block; vertical dashed lines mark the two junctions. Because the mitogenome is circular, reads were mapped to a doubled reference and coordinates folded back onto the 20,520 bp sequence. 29 reads cross the sequence origin and are drawn as two segments on the same row.

## References

Abbott, C. L., Double, M. C., Trueman, J. W. H., Robinson, A., & Cockburn, A. (2005). An unusual source of apparent mitochondrial heteroplasmy: duplicate mitochondrial control regions in *Thalassarche* albatrosses. Molecular Ecology, 14(11), 3605–3613.

Allio, R., Schomaker-Bastos, A., Romiguier, J., Prosdocimi, F., Nabholz, B., & Delsuc, F. (2020). MitoFinder: Efficient automated large-scale extraction of mitogenomic data in target enrichment phylogenomics. Molecular Ecology Resources, 20(4), 892–905.

Astashyn, A., Tvedte, E. S., Sweeney, D., Sapojnikov, V., Bouk, N., Joukov, V., Mozes, E., Strope, P. K., Sylla, P. M., Wagner, L., Bidwell, S. L., Brown, L. C., Clark, K., Davis, E. W., Smith-White, B., Hlavina, W., Pruitt, K. D., Schneider, V. A., & Murphy, T. D. (2024). Rapid and sensitive detection of genome contamination at scale with FCS-GX. Genome Biology, 25(1), 60.

Bao, W., Kojima, K. K., & Kohany, O. (2015). Repbase Update, a database of repetitive elements in eukaryotic genomes. Mobile DNA, 6(1), 11.

Barbraud, C., Delord, K., Bost, C. A., Chaigne, A., Marteau, C., & Weimerskirch, H. (2020). Population trends of penguins in the French Southern Territories. Polar Biology, 43(7), 835–850.

Bardon, G., Barracho, T., Durant, J.M., Lecomte, N., Le Maho, Y., Stenseth, N.C., Cristofari, R. and Le Bohec, C., 2026. Multiannual environmental forcing shapes breeding phenology and success in a sub-Antarctic seabird. Science Advances, 12(11), p.eaea6342.

Bardon, G., Cristofari, R., Winterl, A., Barracho, T., Benoiste, M., Ceresa, C., Chatelain, N., Courtecuisse, J., Fernandes, F. A. N., Gauthier-Clerc, M., Gendner, J.-P., Handrich, Y., Houstin, A., Krellenstein, A., Lecomte, N., Salmon, C.-E., Trucchi, E., Vallas, B., Wong, E. M.,…Le Bohec, C. (2023). RFIDeep: Unfolding the potential of deep learning for radio-frequency identification. Methods in Ecology and Evolution, 14(11), 2814–2826.

Baril, T., Galbraith, J., & Hayward, A. (2024). Earl Grey: a fully automated user-friendly transposable element annotation and analysis pipeline. Molecular Biology and Evolution, 41(4), msae068.

Blum, M., Chang, H.-Y., Chuguransky, S., Grego, T., Kandasaamy, S., Mitchell, A., Nuka, G., Paysan-Lafosse, T., Qureshi, M., Raj, S., Richardson, L., Salazar, G. A., Williams, L., Bork, P., Bridge, A., Gough, J., Haft, D. H., Letunic, I., Marchler-Bauer, A.,…Finn, R. D. (2021). The InterPro protein families and domains database: 20 years on. Nucleic Acids Research, 49(D1), D344–D354.

Boersma, P. D. (2008). Penguins as Marine Sentinels. Bioscience, 58(7), 597–607.

Bosse, M., Spurgin, L. G., Laine, V. N., Cole, E. F., Firth, J. A., Gienapp, P., Gosler, A. G., McMahon, K., Poissant, J., Verhagen, I., Groenen, M. A. M., van Oers, K., Sheldon, B. C., Visser, M. E., & Slate, J. (2017). Recent natural selection causes adaptive evolution of an avian polygenic trait. Science (New York, N.Y.), 358(6361), 365–368.

Böning, C. W., Dispert, A., Visbeck, M., Rintoul, S. R., & Schwarzkopf, F. U. (2008). The response of the Antarctic Circumpolar Current to recent climate change. Nature Geoscience, 1(12), 864–869.

Bost, C. A., Cotté, C., Terray, P., Barbraud, C., Bon, C., Delord, K., Gimenez, O., Handrich, Y., Naito, Y., Guinet, C., & Weimerskirch, H. (2015). Large-scale climatic anomalies affect marine predator foraging behaviour and demography. Nature Communications, 6, 8220.

Brown, M. R., Manuel Gonzalez de La Rosa, P., & Blaxter, M. (2025). Tidk: A toolkit to rapidly identify telomeric repeats from genomic datasets. Bioinformatics (Oxford, England), 41(2), btaf049.

Brůna, T., Lomsadze, A., & Borodovsky, M. (2024). GeneMark-ETP significantly improves the accuracy of automatic annotation of large eukaryotic genomes. Genome Research, 34(5), 757–768.

Buchfink, B., Reuter, K., & Drost, H.-G. (2021). Sensitive protein alignments at tree-of-life scale using DIAMOND. Nature Methods, 18(4), 366–368.

Cantalapiedra, C. P., Hernández-Plaza, A., Letunic, I., Bork, P., & Huerta-Cepas, J. (2021). eggNOG-mapper v2: functional annotation, orthology assignments, and domain prediction at the metagenomic scale. Molecular Biology and Evolution, 38(12), 5825– 5829.

Cardozo, D., Ledesma, M. A., Leotta, G. A., & Coria, N. (2003). El cariotipo del pingüino Rey y del pingüino Emperador (Aves: Sphenisciformes). Journal of Basic & Applied Genetics XV (Suppl. 2), 104.

Chan, P. P., Lin, B. Y., Mak, A. J., & Lowe, T. M. (2021). tRNAscan-SE 2.0: improved detection and functional classification of transfer RNA genes. Nucleic Acids Research, 49(16), 9077–9096.

Chen, S., Zhou, Y., Chen, Y., & Gu, J. (2018). fastp: an ultra-fast all-in-one FASTQ preprocessor. Bioinformatics, 34(17), i884–i890.

Cheng, H., Concepcion, G. T., Feng, X., Zhang, H., & Li, H. (2021). Haplotype-resolved de novo assembly using phased assembly graphs with hifiasm. Nature Methods, 18(2), 170–175.

Cherel, Y., Gilles, J., Handrich, Y., & Le Maho, Y. (1994). Nutrient reserve dynamics and energetics during long-term fasting in the king penguin (*Aptenodytes patagonicus*). Journal of Zoology, 234(1), 1–12.

Clessin, A., Briand, F.-X., Tornos, J., Lejeune, M., De Pasquale, C., Fischer, R., Souchaud, F., Hirchaud, E., Bralet, T., Guinet, C., McMahon, C. R., Grasland, B., Baele, G., & Boulinier, T. (2025). Mass mortality events in the sub-Antarctic Indian Ocean caused by long-distance circumpolar spread of highly pathogenic avian influenza H5N1 clade 2.3.4.4b. bioRxiv: 10.1101/2025.02.25.640068

Cole, T. L., Zhou, C., Fang, M., Pan, H., Ksepka, D. T., Fiddaman, S. R., Emerling, C. A., Thomas, D. B., Bi, X., Fang, Q., Ellegaard, M. R., Feng, S., Smith, A. L., Heath, T. A., Tennyson, A. J. D., Borboroglu, P. G., Wood, J. R., Hadden, P. W., Grosser, S.,…Zhang, G. (2022). Genomic insights into the secondary aquatic transition of penguins. Nature Communications, 13(1), 3912.

Cristofari, R., Davis, L.R., Bardon, G. Nitta Fernandes, F.A., Figueroa, M.E., Franzenburg, S., Gauthier-Clerc, M., Grande, F., Heidrich, R., Hukkanen, M. and Le Maho, Y., Ollikainen, M., Paciello, E., Rampal, P., Stenseth, N.C., Trucchi, E., Zahn, S., Le Bohec, C., Meyer, B.S. (2026). Lifestyle change accelerates epigenetic ageing in King penguins. Nat Commun 17, 3795.

Cristofari, R., Liu, X., Bonadonna, F., Cherel, Y., Pistorius, P., Le Maho, Y., Raybaud, V., Stenseth, N.C., Le Bohec, C. & Trucchi, E. (2018). Climate-driven range shifts of the king penguin in a fragmented ecosystem. Nature Climate Change, 8(3), 245–251.

Dainat, J. (2024). AGAT: Another Gff Analysis Toolkit to handle annotations in any GTF/GFF format. (Version v1.4.1). https://zenodo.org/records/13799920

Danecek, P., Bonfield, J. K., Liddle, J., Marshall, J., Ohan, V., Pollard, M. O., Whitwham, A., Keane, T., McCarthy, S. A., Davies, R. M., & Li, H. (2021). Twelve years of SAMtools and BCFtools. GigaScience, 10(2), giab008.

Descamps, S., Gauthier-Clerc, M., Gendner, J.-P., & Le Maho, Y. (2002). The annual breeding cycle of unbanded king penguins *Aptenodytes patagonicus* on Possession Island (Crozet). Avian Science, 2(2), 87–98.

Dierckxsens, N., Mardulyn, P., & Smits, G. (2017). NOVOPlasty: de novo assembly of organelle genomes from whole genome data. Nucleic Acids Research, 45(4), e18.

Dobin, A., Davis, C. A., Schlesinger, F., Drenkow, J., Zaleski, C., Jha, S., Batut, P., Chaisson, M., & Gingeras, T. R. (2013). STAR: ultrafast universal RNA-seq aligner. Bioinformatics, 29(1), 15–21.

Du, J., Tian, J.-S., Lu, Z.-C., Zhang, S.-J., Song, X.-R., Liu, G.-Y., & Han, J.-B. (2019). Identification of the complete mitochondrial genome of the king penguin *Aptenodytes patagonicus* (Sphenisciformes: Spheniscidae: Aptenodytes). Mitochondrial DNA. Part B, Resources, 4(2), 2191–2192.

Ewels, P.A., Peltzer, A., Fillinger, S. Patel, H., Alneberg, J., Wilm, A., Garcia, M.U., Di Tommaso, P. & Nahnsen, S. (2020). The nf-core framework for community-curated bioinformatics pipelines. Nat Biotechnol 38, 276–278.

Farrell, C., Thompson, M., Tosevska, A., Oyetunde, A., & Pellegrini, M. (2021). BiSulfite Bolt: A bisulfite sequencing analysis platform. GigaScience, 10(5), giab033.

Flynn, J. M., Hubley, R., Goubert, C., Rosen, J., Clark, A. G., Feschotte, C., & Smit, A. F. (2020). RepeatModeler2 for automated genomic discovery of transposable element families. Proceedings of the National Academy of Sciences of the United States of America, 117(17), 9451–9457.

Formenti, G., Rhie, A., Balacco, J., Haase, B., Mountcastle, J., Fedrigo, O., Brown, S., Capodiferro, M. R., Al-Ajli, F. O., Ambrosini, R., Houde, P., Koren, S., Oliver, K., Smith, M., Skelton, J., Betteridge, E., Dolucan, J., Corton, C., Bista, I.,…Vertebrate Genomes Project Consortium. (2021). Complete vertebrate mitogenomes reveal widespread repeats and gene duplications. Genome Biology, 22(1), 120.

Gabriel, L., Brůna, T., Hoff, K. J., Ebel, M., Lomsadze, A., Borodovsky, M., & Stanke, M. (2024). BRAKER3: Fully automated genome annotation using RNA-seq and protein evidence with GeneMark-ETP, AUGUSTUS, and TSEBRA. Genome Research, 34(5), 769–777.

Gabriel, L., Hoff, K. J., Brůna, T., Borodovsky, M., & Stanke, M. (2021). TSEBRA: transcript selector for BRAKER. BMC Bioinformatics, 22(1), 566.

Gherardi-Fuentes, C., Pütz, K., Anguita, C., & Simeone, A. (2019). Comparative foraging and diving behaviour of coexisting breeding and non-breeding King Penguins (*Aptenodytes patagonicus*) in Tierra del Fuego, Chile. The Emu, 119(1), 61–70.

Gunski, R. J., Cañedo, A. D., Garnero, A. D. V., Ledesma, M. A., Coria, N., Montalti, D., & Degrandi, T. M. (2017). Multiple sex chromosome system in penguins (Pygoscelis, Spheniscidae). Comparative Cytogenetics, 11(3), 541–552.

Gurevich, A., Saveliev, V., Vyahhi, N., & Tesler, G. (2013). QUAST: quality assessment tool for genome assemblies. Bioinformatics, 29(8), 1072–1075.

Harrison, P. W., Amode, M. R., Austine-Orimoloye, O., Azov, A. G., Barba, M., Barnes, I., Becker, A., Bennett, R., Berry, A., Bhai, J., Bhurji, S. K., Boddu, S., Branco Lins, P. R., Brooks, L., Ramaraju, S. B., Campbell, L. I., Martinez, M. C., Charkhchi, M., Chougule, K.,…Yates, A. D. (2024). Ensembl 2024. Nucleic Acids Research, 52(D1), D891–D899.

Howe, K., Chow, W., Collins, J., Pelan, S., Pointon, D.-L., Sims, Y., Torrance, J., Tracey, A., & Wood, J. (2021). Significantly improving the quality of genome assemblies through curation. GigaScience, 10(1), giaa153.

Huang, N., & Li, H. (2023). compleasm: a faster and more accurate reimplementation of BUSCO. Bioinformatics, 39(10), btad595.

Huang, Z., Xu, Z., Bai, H., Huang, Y., Kang, N., Ding, X., Liu, J., Luo, H., Yang, C., Chen, W., Guo, Q., Xue, L., Zhang, X., Xu, L., Chen, M., Fu, H., Chen, Y., Yue, Z., Fukagawa, T.,…Xu, L. (2023). Evolutionary analysis of a complete chicken genome. Proceedings of the National Academy of Sciences of the United States of America, 120(8), e2216641120.

Huerta-Cepas, J., Szklarczyk, D., Heller, D., Hernández-Plaza, A., Forslund, S. K., Cook, H., Mende, D. R., Letunic, I., Rattei, T., Jensen, L. J., von Mering, C., & Bork, P. (2019). eggNOG 5.0: a hierarchical, functionally and phylogenetically annotated orthology resource based on 5090 organisms and 2502 viruses. Nucleic Acids Research, 47(D1), D309–D314.

Huisman, J., Kruuk, L. E. B., Ellis, P. A., Clutton-Brock, T., & Pemberton, J. M. (2016). Inbreeding depression across the lifespan in a wild mammal population. Proceedings of the National Academy of Sciences of the United States of America, 113(13), 3585–3590.

Hukkanen, M., Jarman, S., Budd, A., Nitta Fernandes, F. A., Ambrosini, R., Anderson, C., Bardon, G., Berry, O., Bitton, P.-P., Bugnyar, T., Caprioli, M., Carlile, N., Cecere, J. G., Cossin-Sevrin, N., Costanzo, A., Corregidor-Castro, A., Davis, L. R., van Dijk, E., Elsner, M.,…Cristofari, R. (2026). The BEAC, an epigenetic clock for birds. In bioRxiv, 10.64898/2026.08.14.744821

Jones, P., Binns, D., Chang, H.-Y., Fraser, M., Li, W., McAnulla, C., McWilliam, H., Maslen, J., Mitchell, A., Nuka, G., Pesseat, S., Quinn, A. F., Sangrador-Vegas, A., Scheremetjew, M., Yong, S.-Y., Lopez, R., & Hunter, S. (2014). InterProScan 5: genome-scale protein function classification. Bioinformatics, 30(9), 1236–1240.

Kalvari, I., Nawrocki, E. P., Argasinska, J., Quinones-Olvera, N., Finn, R. D., Bateman, A., & Petrov, A. I. (2018). Non-coding RNA analysis using the Rfam database. Current Protocols in Bioinformatics, 62(1), e51.

Kalvari, I., Nawrocki, E. P., Ontiveros-Palacios, N., Argasinska, J., Lamkiewicz, K., Marz, M., Griffiths-Jones, S., Toffano-Nioche, C., Gautheret, D., Weinberg, Z., Rivas, E., Eddy, S. R., Finn, R. D., Bateman, A., & Petrov, A. I. (2021). Rfam 14: expanded coverage of metagenomic, viral and microRNA families. Nucleic Acids Research, 49(D1), D192–D200.

Kang, Y.-J., Yang, D.-C., Kong, L., Hou, M., Meng, Y.-Q., Wei, L., & Gao, G. (2017). CPC2: a fast and accurate coding potential calculator based on sequence intrinsic features. Nucleic Acids Research, 45(W1), W12–W16.

Katoh, K., & Standley, D. M. (2013). MAFFT multiple sequence alignment software version 7: improvements in performance and usability. Molecular Biology and Evolution, 30(4), 772–780.

Kim, D., Paggi, J. M., Park, C., Bennett, C., & Salzberg, S. L. (2019). Graph-based genome alignment and genotyping with HISAT2 and HISAT-genotype. Nature Biotechnology, 37(8), 907–915.

Kovaka, S., Zimin, A. V., Pertea, G. M., Razaghi, R., Salzberg, S. L., & Pertea, M. (2019). Transcriptome assembly from long-read RNA-seq alignments with StringTie2. Genome Biology, 20(1), 278.

Kriventseva, E. V., Kuznetsov, D., Tegenfeldt, F., Manni, M., Dias, R., Simão, F. A., & Zdobnov, E. M. (2019). OrthoDB v10: sampling the diversity of animal, plant, fungal, protist, bacterial and viral genomes for evolutionary and functional annotations of orthologs. Nucleic Acids Research, 47(D1), D807–D811.

Krzywinski, M., Schein, J., Birol, I., Connors, J., Gascoyne, R., Horsman, D., Jones, S. J., & Marra, M. A. (2009). Circos: an information aesthetic for comparative genomics. Genome Research, 19(9), 1639–1645.

Kuznetsov, D., Tegenfeldt, F., Manni, M., Seppey, M., Berkeley, M., Kriventseva, E. V., & Zdobnov, E. M. (2023). OrthoDB v11: annotation of orthologs in the widest sampling of organismal diversity. Nucleic Acids Research, 51(D1), D445–D451.

Lagesen, K., Hallin, P., Rødland, E. A., Staerfeldt, H.-H., Rognes, T., & Ussery, D. W. (2007). RNAmmer: consistent and rapid annotation of ribosomal RNA genes. Nucleic Acids Research, 35(9), 3100–3108.

Le Bohec, C., Gauthier-Clerc, M., Grémillet, D., Pradel, R., Béchet, A., Gendner, J.-P., & Le Maho, Y. (2007). Population dynamics in a long-lived seabird: I. Impact of breeding activity on survival and breeding probability in unbanded king penguins. Journal of Animal Ecology, 76(6), 1149–1160.

Le Bohec, C. L., Durant, J. M., Gauthier-Clerc, M., Stenseth, N. C., Park, Y.-H., Pradel, R., Grémillet, D., Gendner, J.-P., & Le Maho, Y. (2008). King penguin population threatened by Southern Ocean warming. Proceedings of the National Academy of Sciences, 105(7), 2493–2497.

Le Vaillant, M., Ropert-Coudert, Y., Le Maho, Y., & Le Bohec, C. (2016). Individual parameters shape foraging activity in breeding king penguins. Behavioral Ecology, 27(1), 352–362.

Le Vaillant, M., Wilson, R. P., Kato, A., Saraux, C., Hanuise, N., Prud’homme, O., Le Maho, Y., Le Bohec, C., & Ropert-Coudert, Y. (2012). King penguins adjust their diving behaviour with age. Journal of Experimental Biology, 215(Pt 21), 3685–3692.

Lewin, H. A., Robinson, G. E., Kress, W. J., Baker, W. J., Coddington, J., Crandall, K. A., Durbin, R., Edwards, S. V., Forest, F., Gilbert, M. T. P., Goldstein, M. M., Grigoriev, I. V., Hackett, K. J., Haussler, D., Jarvis, E. D., Johnson, W. E., Patrinos, A., Richards, S., Castilla-Rubio, J. C.,…Zhang, G. (2018). Earth BioGenome Project: Sequencing life for the future of life. Proceedings of the National Academy of Sciences of the United States of America, 115(17), 4325–4333.

Li, A., Zhang, J., & Zhou, Z. (2014). PLEK: a tool for predicting long non-coding RNAs and messenger RNAs based on an improved k-mer scheme. BMC Bioinformatics, 15(1), 311.

Li, H. (2018). Minimap2: pairwise alignment for nucleotide sequences. Bioinformatics, 34(18), 3094–3100.

Li, H. (2023). Protein-to-genome alignment with miniprot. Bioinformatics, 39(1), btad014.

Love, M. I., Huber, W., & Anders, S. (2014). Moderated estimation of fold change and dispersion for RNA-seq data with DESeq2. Genome Biology, 15(12), 550.

Manni, M., Berkeley, M. R., Seppey, M., Simão, F. A., & Zdobnov, E. M. (2021). BUSCO update: novel and streamlined workflows along with broader and deeper phylogenetic coverage for scoring of eukaryotic, prokaryotic, and viral genomes. Molecular Biology and Evolution, 38(10), 4647–4654.

Marçais, G., Delcher, A. L., Phillippy, A. M., Coston, R., Salzberg, S. L., & Zimin, A. (2018). MUMmer4: A fast and versatile genome alignment system. PLoS Computational Biology, 14(1), e1005944.

Mathers, T.C., Paulini, M., Sotero-Caio, C.G., & Wood J.M.D. (2026). MicroFinder: Conserved gene-set mapping and assembly ordering for manual curation of bird microchromosomes. GigaScience, 15, giag036.

Mikheenko, A., Prjibelski, A., Saveliev, V., Antipov, D., & Gurevich, A. (2018). Versatile genome assembly evaluation with QUAST-LG. Bioinformatics, 34(13), i142–i150.

Montaña-Lozano, P., Moreno-Carmona, M., Ochoa-Capera, M., Medina, N. S., Boore, J. L., & Prada, C. F. (2022). Comparative genomic analysis of vertebrate mitochondrial reveals a differential of rearrangements rate between taxonomic class. Scientific Reports, 12(1), 5479.

Morris-Pocock, J. A., Taylor, S. A., Birt, T. P., & Friesen, V. L. (2010). Concerted evolution of duplicated mitochondrial control regions in three related seabird species. BMC Evolutionary Biology, 10(1), 14.

Nawrocki, E. P., & Eddy, S. R. (2013). Infernal 1.1: 100-fold faster RNA homology searches. Bioinformatics, 29(22), 2933–2935.

Nevers, Y., Warwick Vesztrocy, A., Rossier, V., Train, C.-M., Altenhoff, A., Dessimoz, C., & Glover, N. M. (2024). Quality assessment of gene repertoire annotations with OMArk. Nature Biotechnology, 43(1), 124–133.

Nitta Fernandes, F.A., Bardon, G., Paris. J.R., Cristofari, R., Ferrer Obiol, J., Ancona, L., Greco, S., Gerdol, M., Vallas, B., Carette, P., Le Bohec, C., & Trucchi, E. (2025). A long-lived seabird buffers the cost of phenological mismatch by modulating stress response, bioRxiv: 10.1101/2025.07.10.663151.

Oberlin, M., Enstipp, M.R., Le Bohec, C., Dardel, R., Bost, C.-A., Handrich, Y. (2025). Exploring new depths: king penguins break dive records during the austral winter. J Exp Biol 1, 228(7): jeb250184.

Olsson, O. (1996). Seasonal effects of timing and reproduction in the king penguin: a unique breeding cycle. Journal of Avian Biology, 27(1), 7–14.

Ontiveros-Palacios, N., Cooke, E., Nawrocki, E. P., Triebel, S., Marz, M., Rivas, E., Griffiths-Jones, S., Petrov, A. I., Bateman, A., & Sweeney, B. (2025). Rfam 15: RNA families database in 2025. Nucleic Acids Research, 53(D1), D258–D267.

Ottenburghs, J., Geng, K., Suh, A., & Kutter, C. (2021). Genome size reduction and transposon activity impact tRNA gene diversity while ensuring translational stability in birds. Genome Biology and Evolution, 13(4), evab016.

Pan, H., Cole, T. L., Bi, X., Fang, M., Zhou, C., Yang, Z., Ksepka, D. T., Hart, T., Bouzat, J. L., Argilla, L. S., Bertelsen, M. F., Boersma, P. D., Bost, C.-A., Cherel, Y., Dann, P., Fiddaman, S. R., Howard, P., Labuschagne, K., Mattern, T.,…Zhang, G. (2019). High-coverage genomes to elucidate the evolution of penguins. GigaScience, 8(9), giz117.

Paris, J. R., Nitta Fernandes, F. A., Pirri, F., Greco, S., Gerdol, M., Pallavicini, A., Benoiste, M., Cornec, C., Zane, L., Haas, B., Le Bohec, C., & Trucchi, E. (2025). Gene expression shifts in Emperor penguin adaptation to the extreme Antarctic environment. Molecular Ecology, e17552.

Paris JR, Nitta Fernandes FA, Santos CA, Pointon D-LB, Wood JMD, Ferrer Obiol J, Salces-Ortiz J, Fernández R, Cristofari R, Le Bohec C, Trucchi E (2026). King-penguin-genome-assembly. Zenodo. 10.5281/zenodo.22968008

Peona, V., Blom, M. P. K., Frankl-Vilches, C., Milá, B., Ashari, H., Thébaud, C., Benz, B. W., Christidis, L., Gahr, M., Irestedt, M., & Suh, A. (2022). The hidden structural variability in avian genomes. bioRxiv: 10.1101/2021.12.31.473444

Peona, V., Palacios-Gimenez, O. M., Blommaert, J., Liu, J., Haryoko, T., Jønsson, K. A., Irestedt, M., Zhou, Q., Jern, P., & Suh, A. (2021). The avian W chromosome is a refugium for endogenous retroviruses with likely effects on female-biased mutational load and genetic incompatibilities. Philosophical Transactions of the Royal Society of London. Series B, Biological Sciences, 376(1833), 20200186.

Pistorius, P. A., Baylis, A., Crofts, S., & Pütz, K. (2012). Population development and historical occurrence of king penguins at the Falkland Islands. Antarctic Science, 24(5), 435–440.

Pointon, D.-L. B. (2025). sanger-tol/curationpretext. Zenodo. 10.5281/zenodo.12773958 (Version 1.0.1).

Pointon, D.-L. B, Sims, Y., Aunin, E., Eagles, W., Torrance J., Downie, J. (2025). sanger-tol/ascc: v0.1.0 -Red Book (0.1.0). Zenodo. 10.5281/zenodo.14883766 (Version 0.1.0).

Putri, G. H., Anders, S., Pyl, P. T., Pimanda, J. E., & Zanini, F. (2022). Analysing high-throughput sequencing data in Python with HTSeq 2.0. Bioinformatics, 38(10), 2943– 2945.

Ramos, B., González-Acuña, D., Loyola, D. E., Johnson, W. E., Parker, P. G., Massaro, M., Dantas, G. P. M., Miranda, M. D., & Vianna, J. A. (2018). Landscape genomics: natural selection drives the evolution of mitogenome in penguins. BMC Genomics, 19(1), 53.

Ranallo-Benavidez, T. R., Jaron, K. S., & Schatz, M. C. (2020). GenomeScope 2.0 and Smudgeplot for reference-free profiling of polyploid genomes. Nature Communications, 11(1), 1–10.

Rhie, A., McCarthy, S. A., Fedrigo, O., Damas, J., Formenti, G., Koren, S., Uliano-Silva, M., Chow, W., Fungtammasan, A., Kim, J., Lee, C., Ko, B. J., Chaisson, M., Gedman, G. L., Cantin, L. J., Thibaud-Nissen, F., Haggerty, L., Bista, I., Smith, M.,…Jarvis, E. D. (2021). Towards complete and error-free genome assemblies of all vertebrate species. Nature, 592(7856), 737–746.

Rhie, A., Walenz, B. P., Koren, S., & Phillippy, A. M. (2020). Merqury: reference-free quality, completeness, and phasing assessment for genome assemblies. Genome Biology, 21(1), 245.

Rizk, G., Lavenier, D., & Chikhi, R. (2013). DSK: k-mer counting with very low memory usage. Bioinformatics, 29(5), 652–653.

Gimeno, M., Giménez, J., Chiaradia, A., Davis, L. S., Seddon, P. J., Ropert-Coudert, Y., Reisinger, R. R., Coll, M., & Ramírez, F. (2024). Climate and human stressors on global penguin hotspots: Current assessments for future conservation. Global Change Biology, 30(1), e17143.

Sangster, G., & Luksenburg, J. A. (2021). Sharp increase of problematic mitogenomes of birds: Causes, consequences, and remedies. Genome Biology and Evolution, 13(9), evab210.

Saraux, C., Friess, B., Le Maho, Y., & Le Bohec, C. (2012). Chick-provisioning strategies used by king penguins to adapt to a multiseasonal breeding cycle. Animal Behaviour, 84(3), 675–683.

Scheffer, A., Trathan, P. N., & Collins, M. (2010). Foraging behaviour of King Penguins (*Aptenodytes patagonicus*) in relation to predictable mesoscale oceanographic features in the Polar Front Zone to the north of South Georgia. Progress in Oceanography, 86(1-2), 232–245.

Smit, A. F. A., Hubley, R., & Green, P. (2015). RepeatMasker Open-4.0. 2013-2015. http://www.repeatmasker.org

Sozzoni, M., Ferrer Obiol, J., Formenti, G., Tigano, A., Paris, J. R., Balacco, J. R.,…& Rubolini, D. (2023). A chromosome-level reference genome for the Black-Legged Kittiwake (*Rissa tridactyla*), a declining circumpolar seabird. Genome Biology and Evolution, 15(8), evad153.

Stanke, M., Diekhans, M., Baertsch, R., & Haussler, D. (2008). Using native and syntenically mapped cDNA alignments to improve de novo gene finding. Bioinformatics, 24(5), 637–644.

Stanke, M., Schöffmann, O., Morgenstern, B., & Waack, S. (2006). Gene prediction in eukaryotes with a generalized hidden Markov model that uses hints from external sources. BMC Bioinformatics, 7, 62.

Stier, A., Viblanc, V. A., Massemin-Challet, S., Handrich, Y., Zahn, S., Rojas, E. R., Saraux, C., Le Vaillant, M., Prud’homme, O., Grosbellet, E., Robin, J.-P., Bize, P., & Criscuolo, F. (2014). Starting with a handicap: phenotypic differences between early and late born king penguin chicks and their survival correlates. Functional Ecology, 28(3), 601–611.

Stoffel, M. A., Johnston, S. E., Pilkington, J. G., & Pemberton, J. M. (2021). Genetic architecture and lifetime dynamics of inbreeding depression in a wild mammal. Nature Communications, 12(1), 2972.

Stonehouse, B. (1960) The king penguin *Aptenodytes patagonicus* of South Georgia I. Breeding behavior and development. Sci Rep Falkl Isl Depend Surv., 23:1–81.

Storer, J., Hubley, R., Rosen, J., Wheeler, T. J., & Smit, A. F. (2021). The Dfam community resource of transposable element families, sequence models, and genome annotations. Mobile DNA, 12(1), 2.

Tang, S., Lomsadze, A., & Borodovsky, M. (2015). Identification of protein coding regions in RNA transcripts. Nucleic Acids Research, 43(12), e78.

Thorvaldsdóttir, H., Robinson, J. T., & Mesirov, J. P. (2013). Integrative Genomics Viewer (IGV): high-performance genomics data visualization and exploration. Briefings in Bioinformatics, 14(2), 178–192.

Trucchi, E., Gratton, P., Whittington, J. D., Cristofari, R., Le Maho, Y., Stenseth, N. C., & Le Bohec, C. (2014). King penguin demography since the last glaciation inferred from genome-wide data. Proceedings of the Royal Society B: Biological Sciences, 281(1787), 20140528.

Trucchi, E., Massa, P., Giannelli, F., Latrille, T., Gargano M., Nitta Fernandes, F. A., Ancona, L., Stenseth, N. C., Ferrer Obiol, J., Paris, J. R., Bertorelle, G., & Le Bohec, C. (2025). High gene expression predicts extremely low segregation of deleterious mutations in large penguin populations. Molecular Biology and Evolution, 42(6), msaf146.

Uliano-Silva, M., Ferreira, J. G. R. N., Krasheninnikova, K., Darwin Tree of Life Consortium, Formenti, G., Abueg, L., Torrance, J., Myers, E. W., Durbin, R., Blaxter, M., & McCarthy, S. A. (2023). MitoHiFi: a python pipeline for mitochondrial genome assembly from PacBio high fidelity reads. BMC Bioinformatics, 24(1), 288.

UniProt Consortium. (2025). UniProt: The universal protein knowledgebase in 2025. Nucleic Acids Research, 53(D1), D609–D617.

Urantówka, A. D., Kroczak, A., & Mackiewicz, P. (2020). New view on the organization and evolution of Palaeognathae mitogenomes poses the question on the ancestral gene rearrangement in Aves. BMC Genomics, 21(1), 874.

Vaisvila, R., Ponnaluri, V. K. C., Sun, Z., Langhorst, B. W., Saleh, L., Guan, S., Dai, N., Campbell, M. A., Sexton, B. S., Marks, K., Samaranayake, M., Samuelson, J. C., Church, H. E., Tamanaha, E., Corrêa, I. R., Jr, Pradhan, S., Dimalanta, E. T., Evans, T. C., Jr, Williams, L., & Davis, T. B. (2021). Enzymatic methyl sequencing detects DNA methylation at single-base resolution from picograms of DNA. Genome Research, 31(7), 1280–1289.

Van Heezik, Y.M., Seddon, P.J., Cooper, J. and Plös, A.L. (1994). Interrelationships between breeding frequency, timing and outcome in King Penguins *Aptenodytes patagonicus*: are King Penguins biennial breeders? Ibis, 136: 279–284.

Wang, L., Park, H. J., Dasari, S., Wang, S., Kocher, J.-P., & Li, W. (2013). CPAT: Coding-Potential Assessment Tool using an alignment-free logistic regression model. Nucleic Acids Research, 41(6), e74.

Waters, P. D., Patel, H. R., Ruiz-Herrera, A., Álvarez-González, L., Lister, N. C., Simakov, O., Ezaz, T., Kaur, P., Frere, C., Grützner, F., Georges, A., & Graves, J. A. M. (2021). Microchromosomes are building blocks of bird, reptile, and mammal chromosomes. Proceedings of the National Academy of Sciences of the United States of America, 118(45), e2112494118.

Xu, L., Auer, G., Peona, V., Suh, A., Deng, Y., Feng, S., Zhang, G., Blom, M. P. K., Christidis, L., Prost, S., Irestedt, M., & Zhou, Q. (2019). Dynamic evolutionary history and gene content of sex chromosomes across diverse songbirds. Nature Ecology & Evolution, 3(5), 834–844.

Zhang, H., Song, L., Wang, X., Cheng, H., Wang, C., Meyer, C. A., Liu, T., Tang, M., Aluru, S., Yue, F., Liu, X. S., & Li, H. (2021). Fast alignment and preprocessing of chromatin profiles with Chromap. Nature Communications, 12(1), 6566.

Zhou, C., McCarthy, S. A., & Durbin, R. (2023). YaHS: yet another Hi-C scaffolding tool. Bioinformatics, 39(1), btac808.

